# Phylogeny inference under the general Markov model using MST-backbone

**DOI:** 10.1101/2020.06.30.180315

**Authors:** Prabhav Kalaghatgi

## Abstract

**Motivation:** Phylogeny inference via maximum likelihood is NP-hard. Current methods make simplifying assumptions such as stationarity, homogeneity, and time-reversibility for computational ease. The stationarity assumption is violated by empirical observations of GC content evolution, and might systematically bias phylogeny inference. The general Markov model (GM) is a suitable alternative to stationary models because it allows for the evolution of GC content. Related work on the GM model has predominantly focused on inferring unrooted trees using either the log-det distance or phylogenetic invariants.

**Methods:** We adapted the structural EM framework to perform tree search under the GM model (SEM-GM). Additionally, we implemented a minimum spanning tree framework called MST-backbone to improve the scalability of SEM-GM by constraining search through tree space. MST-backbone(SEM-GM) was used to infer unrooted trees, which are subsequently rooted under the GM model; the latter procedure is called rSEM-GM. We compared our method with RAxML-NG, IQ-TREE, and FastTree on simulated data. We validated our methods on six empirical datasets.

**Results:** Estimated experimental phylogenies are rooted with high accuracy under the GM model (recall ranging from 80% to 94%). However, virus phylogenies are not realistically rooted, suggesting that the GM model may be overtrained on some empirical datasets. The comparative analysis of simulated data suggests that MST-backbone(SEM-GM) and FastTree scale linearly whereas rSEM-GM, RAxML-NG, and IQ-TREE scale quadratically. The results on empirical data suggest that it is not necessary to use the general time-reversible model for computational ease.

**Availability:** https://github.com/prabhavk/mst-backbone-sem-gm

**Supplementary information:** Supplementary data are available online

## 1 Introduction

Probabilistic models are widely used to model gene evolution, estimate molecular substitution rates, and infer evolutionary history. The probabilistic models that are commonly used are continuous-time hidden Markov models (CT-HMM) on leaf-labeled phylogenetic trees. The optimization problem of finding maximum likelihood (ML) phylogenies is NP-hard, and the corresponding decision problem is NP-complete (Chickering, 1996; Chor and Tuller, 2006; Roch, 2006). The majority of widely used software performs extensive search through tree space, either via sampling procedures (Hohna *et al.*, 2016; Bouckaert *et al.*, 2014), or by tree modification operations (Minh *et al.*, 2020; Kozlov *et al.*, 2019). CT-HMM are parameterized in terms of rate matrices, branch lengths, and a probability distribution for the hidden sequence at the root. Current software makes simplifying assumptions about the underlying Markov process such as stationarity, homogeneity and time-reversibility in order to exhaustively search through tree space.

The stationarity assumption is violated by empirical observations of evolution in GC content (Agashe and Shankar, 2014; Nabholz *et al.*, 2011). Spontaneous deamination of methylated cytosine, mutation bias and GC biased gene conversion are some of the molecular mechanisms underlying evolution of GC content (Ehrlich *et al.*, 1990; Mugal *et al.*, 2015). The widespread use of time-reversible models is thought to have systematically biased phylogeny reconstruction by grouping together sequences based on similarity in GC content (Betancur-R *et al.*, 2013). Sheffield *et al.* (2009) found that trees inferred using time-reversible models were not compatible with established taxonomic relationships, whereas trees inferred using non-stationary CT-HMMs (specifically, p4 by Foster (2004) and PHASE by Gowri-Shankar and Rattray (2007)) were compatible with established taxonomic relationships.

The general Markov model (GM) by Barry and Hartigan (1987) is a non-stationary, non-homogeneous, and time-irreversible discrete-time hidden Markov model (DT-HMM) on trees that is suitable for modeling the evolution of GC content. The GM model is parameterized in terms of a probability distribution for the hidden sequence at the root, and transition matrices for each edge of the tree under consideration. Pachter and Sturmfels (2005) noted that the GM model generalizes the hidden Markov model (HMM), a model that is widely used for finding sequence motifs (Cappé *et al.*, 2005). The HMM can be optimized using an expectation-maximization algorithm (EM; Dempster *et al.* (1977)) known as the Baum-Welch algorithm (Sammut and Webb, 2010). Consequently, it should be possible to optimize the parameters of the GM model using an EM algorithm.

As far as we are aware, the literature on inferring phylogenies using the GM model is limited to inferring unrooted trees. Given sequences generated under the GM model on a rooted tree, it is possible to infer the unrooted topology using distance-based methods such as neighbor-joining (NJ; Saitou and Nei (1987)) using log-det distances (Steel, 1994). Eriksson (2005) and Allman *et al.* (2017) provide methods for constructing unrooted trees by inferring splits using techniques based on phylogenetic invariants.

In this paper, we adapt Friedman’s structural EM algorithm to phylogeny inference under the general Markov model (Friedman, 1997; Friedman *et al.*, 2002). We refer to the new method as SEM-GM. Additionally, we implemented a minimum spanning tree (MST) framework called MST-backbone in order to improve the scalability of SEM-GM by constraining the search through tree space.

The design of MST-backbone is inspired by the topological relationship between unrooted phylogenetic trees and MSTs computed using additive distances that was introduced by Choi *et al.* (2011). Kalaghatgi and Lengauer (2017) introduced a special class of MSTs called vertex-order based MSTs in order to correct the proof of Choi *et al.*. We do not assume that distances are additive in this paper. In recent work, Zhang *et al.* (2019) presented a MST-based method called incremental tree construction (INC) that uses MSTs to select quartets that are subsequently attached to an incrementally growing three-leaf phylogenetic tree. Le *et al.* (2019) implemented a version of INC called INC-ML that builds phylogenetic trees by using trees obtained using RAxML (Stamatakis, 2006) and FastTree (Price *et al.*, 2010). Le *et al.* (2019) found that RAxML outperformed FastTree and INC-ML across a wide range of simulation scenarios.

MST-backbone was designed to construct an unrooted phylogenetic tree. We implemented an EM procedure for rooting an unrooted tree under the GM model called restricted SEM-GM (rSEM-GM). Additionally, we implemented a model selection procedure called UNRESTselector because the GM model has a large number of free parameters and may be prone to overtraining. UNRESTselector roots trees using CT-HMM that are constructed using one or more distinct UNREST rate matrices (Yang, 1994), where the number of distinct rate matrices is selected using Bayesian information criterion (BIC).

In the following text MST-backbone(SEM-GM)+rSEM-GM and MST-backbone(SEM-GM)+UNRESTselector refer to methods that construct unrooted trees using MST-backbone(SEM-GM) and subsequently root the trees under the GM model and the CT-HMM selected by UNRESTselector, respectively.

In the current study we perform a comparative analysis of MST-backbone(SEM-GM) and MST-backbone(SEM-GM)+rSEM-GM with IQ-TREE v2.0 (Minh *et al.*, 2020), RAxML-NG v0.8.1 (Kozlov *et al.*, 2019) and FastTree v2.1.10 (Price *et al.*, 2010) using simulated sequences where GC content was allowed to evolve. We compared MST-backbone(SEM-GM)+rSEM-GM and MST-backbone(SEM-GM) +UNRESTselector with IQ-TREE on six empirical datasets. Twenty empirical alignments were analyzed in total.

## 2 Methods

### 2.1 Hidden Markov models on trees

A phylogenetic tree is a rooted tree with hidden vertices representing unsampled ancestors and labeled leaf vertices representing sampled taxa. The in-degree of each hidden vertex is one except for one vertex that has in-degree zero (the root *ρ*). If the out-degree of each hidden vertex of a phylogenetic tree is two then the phylogenetic tree is said to be a bifurcating phylogenetic tree. Phylogenetic trees are assumed to be bifurcating unless specified otherwise. The set of hidden vertices and the set of labeled vertices are denoted by *H* and *L*, respectively. In work presented here, multiple sequence alignments of DNA sequences are used to infer phylogenetic trees. The variable 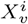 denotes the DNA base at position (site) *i* in the sequence of the vertex *v*. The variables *X_h_*: *h* ∈ *H* are hidden variables, and the variables *X_l_*: *l* ∈ *L* are observed variables.

The general Markov model (GM; Barry and Hartigan (1987)) *M* on a phylogenetic tree *T* = (*V, E*) is a discrete-time hidden Markov model that is parameterized in terms of an independent transition matrix *P*(*u,v*) for each directed edge (*u, v*) in *E*, and a root probability distribution *π_ρ_*. A continuous-time hidden Markov model (CT-HMM) on a phylogenetic tree *T* = (*V, E*) is parameterized in terms of branch lengths {*t_e_*: *e* ∈ *E*}, rate matrices {*Q_e_*: *e* ∈ *E*}, and a root probability distribution *π_ρ_*. The transition matrices for CT-HMM are computed as 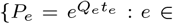 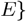.

The restrictions that are commonly placed on CT-HMM are homogeneity, stationarity, and time-reversibility. A CT-HMM is homogeneous if the rate matrices *Q_e_* are identical. A homogeneous CT-HMM is stationary if base composition does not evolve across any edge, a condition that is satisfied if *π_ρ_Q* = 0 (Steel, 2016) where *Q* is the rate matrix that acts on each edge. The UNREST model is a stationary homogeneous CT-HMM. The GTR model is a stationary homogeneous CT-HMM on trees with the additional restriction that the Markov model is time-reversible, i.e.,

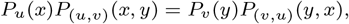

for each edge (*u, v*) in *E*, where *P_u_* is the marginal probability of variable *X_u_* which is identical to the root probability distribution *π_ρ_* for a stationary distribution. Time-reversibility is ensured by constructing a rate matrix *Q* such that Π*Q* is symmetric, where Π is the diagonal matrix such that Π(*i, i*) is the *i*^th^ element of the stationary distribution of the homogeneous Markov model that is parameterized by *Q*.

We describe an expectation maximization procedure for performing tree search under the GM model in the following subsection.

### 2.2 Structural EM for the general Markov model

Expectation-maximization (EM) algorithms are a class of algorithms that are applied to a wide variety of models with hidden variables (Dempster *et al.*, 1977). Structural expectation maximization is an EM algorithm that is designed for Bayesian networks with hidden variables (Friedman, 1997). Structural EM was adapted to phylogeny inference under the GTR model by Friedman *et al.* (2002).

EM algorithms are typically used for fitting models such that parameter estimation can be performed in a computationally efficient manner (typically in closed-form) if there are no hidden variables. EM algorithms simplify the problem of fitting models that contain hidden variables by computing expected values for hidden variables that are sufficient to enable the use of computationally efficient methods to estimate model parameters. Maximum likelihood phylogenetic trees with no hidden variables are known as Chow-Liu trees (Chow and Liu, 1968)). A Chow-Liu tree *T^CL^* can be inferred in polynomial-time by computing a maximum mutual-information spanning tree *T*^MI^ using Prim’s algorithm (Prim, 1957), followed by rooting *T* ^MI^ such that log-likelihood is maximized (Chow and Liu, 1968). The maximum likelihood estimate (MLE) of model parameters of a GM model on *T^CL^* can be computed in closed-form as follows (Koller and Friedman, 2009).

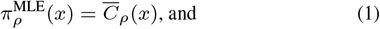

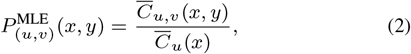

where the observed count matrices 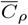 and 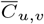 are empirically estimated.

Structural EM for the GM model (SEM-GM) iteratively improves an initial estimate of a GM model on a phylogenetic tree via the expectation step (E-step) and the maximization step (M-step) until the log-likelihood score converges. First, we give an overview of SEM-GM, and then we follow it up with a detailed description. See Fig. 1 for an illustration of SEM-GM.

Given a GM model *M* on a phylogenetic tree *T* = (*V* = {*H* ∪ *L*}, *E*), the E-step involves computing the expected co-occurrence of states *E_M_* [*C_u,v_* (*x, y*)] for each vertex pair *u, v* in *V*, and the expected occurrence of states *E_M_* [*C_h_*(*x*)] for each hidden vertex *h* in *H*. The matrices that are computed in the E-step are referred to as expected count matrices. The M-step uses the expected count matrices to compute a maximum expected likelihood phylogenetic tree. The E-step and the M-step are performed iteratively until improvement in log-likelihood score is smaller than a likelihood convergence threshold. We used a convergence threshold of 0.01 log-likelihood units.

**Fig. 1.**
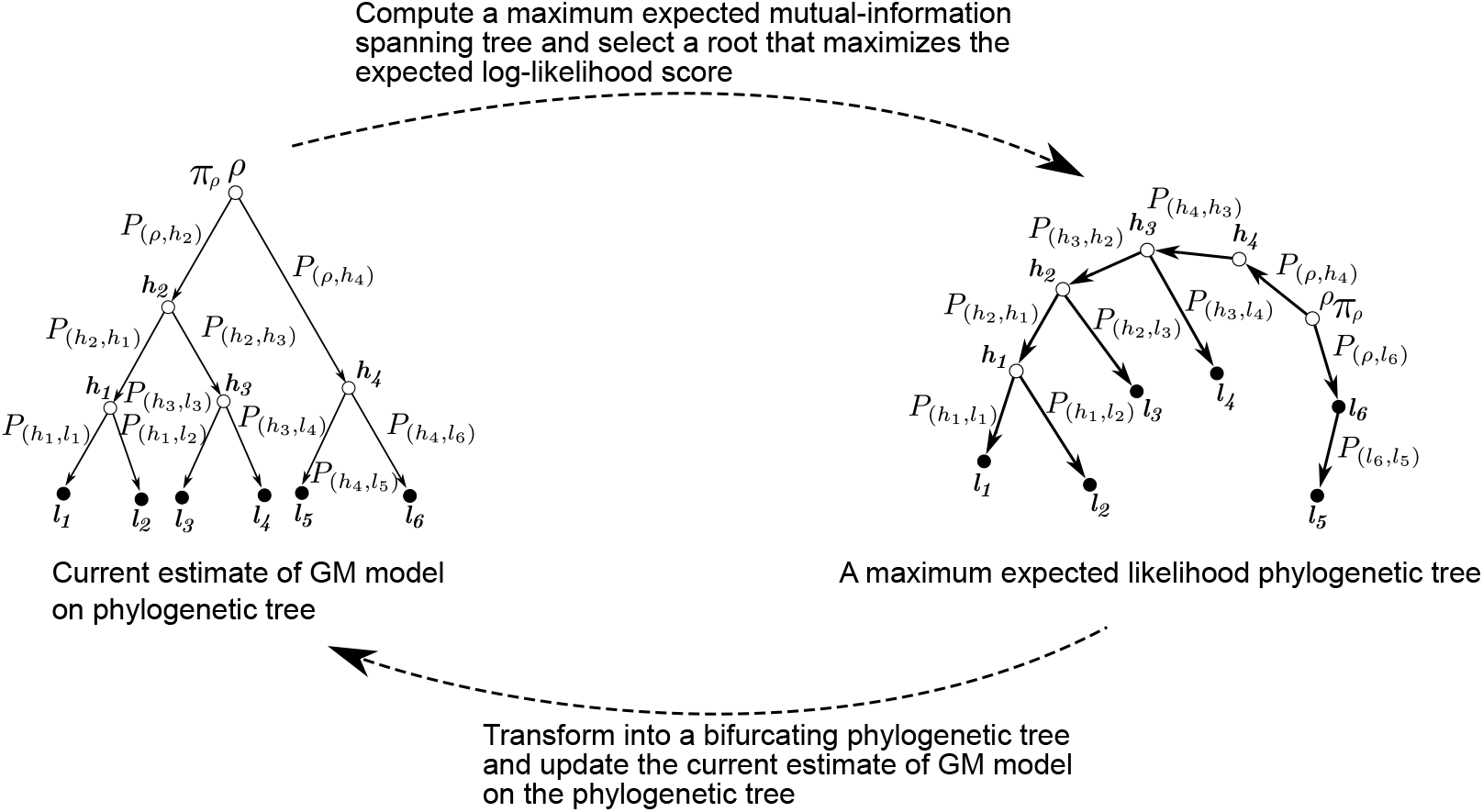
An illustration of structural EM for the general Markov model on phylogenetic trees

#### Computing expected counts

The E-step involves computing expected count matrices, and is described in detail below. The expected count 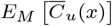 for a hidden vertex *u* is computed as follows (Koller and Friedman, 2009).

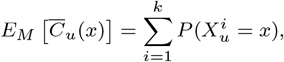

where 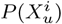 is the marginal probability

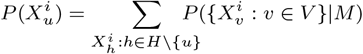

Similarly, the expected count 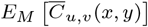 for any vertex pair *u, v* is computed as follows

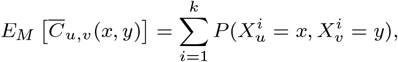

where 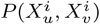 is the marginal probability

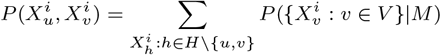

The marginal probability 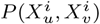 for adjacent vertices *u* and *v* is computed using the belief propagation algorithm (Pearl, 1982; Koller and Friedman, 2009). 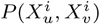 for non-adjacent vertices *u, v* is computed in order of increasing unweighted path length from *u* to *v* on *T* as follows. Consider a path (*v*_1_, *v*_2_,…, *v_n−_*_1_*, v_n_*) in the undirected version of *T* such that 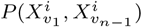 has been computed and we are interested in computing 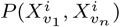. Note that variables 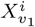 and 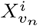 are independent if conditioned on 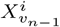. Thus 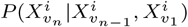 equals 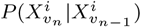. This enables us to decompose 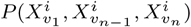 as 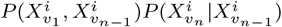.

The marginal probability 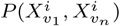 is computed as

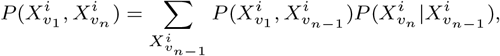

where 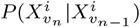 is computed using 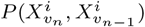.

#### Computing a maximum expected likelihood phylogenetic tree

Expected mutual information scores are computed using marginal probabilities computed in the E-step. A complete graph over hidden vertices and labeled vertices is constructed and the edges are weighted with expected mutual information scores. Subsequently, a maximum expected mutual-information spanning tree *T* ^EMI^ is computed using Prim’s algorithm. Finally, *T* ^EMI^ is rooted to construct *T* ^EL^ such that the expected likelihood score is maximized. The MLE 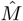 of a GM model on *T* ^MEL^ is computed in closed-form by applying equations (1) and (2) to expected count matrices. Note that *T* ^EL^ might not be a bifurcating phylogenetic tree and may contain non-leaf labeled vertices. *T* ^EL^ is transformed if *T* ^EL^ is not a bifurcating phylogenetic tree by swapping edges as necessary and modifying the estimated GM model such that the log-likelihood score remains unchanged after the transformation operations. The transformation operations are described in detail in Section 1 of the supplementary material.

#### Computing an initial estimate for SEM-GM

SEM-GM is initialized by (*i*) constructing an unrooted phylogenetic tree using neighbor-joining (NJ; Saitou and Nei (1987)), (*ii*) rooting the unrooted tree by inserting a hidden vertex along an edge that is selected at random, (*iii*) inferring sequences for each hidden vertex via maximum parsimony using Fitch-Hartigan algorithm (Fitch, 1971; Hartigan, 1973), and (*iv*) estimating the parameters of a GM model using equations (1) and (2), where the count matrices are constructed using the observed sequences and the sequences for hidden vertices that were inferred in step (*iii*).

### 2.3 MST-backbone

SEM-GM is computationally expensive because each E-step involves *O*(*n*^2^*a*^3^*k*) operations, where *n* is the number of vertices in *V*, *a* is the size of the alphabet (four for DNA), and *k* is the number of alignment columns. We developed a minimum spanning tree framework called MST-backbone to constrain the search through tree space in order to improve the scalability of SEM-GM.

MST-backbone builds a global unrooted phylogenetic tree T in an agglomerative fashion. The main steps of MST-backbone are (*i*) computing a minimum spanning tree (MST) using Hamming distances; (*ii*) selecting smallest mutually independent vertex sets *V_s_* and *V_e_* comprising at least *s* vertices each such that (*a*) *V_s_* induces a subtree in *M*, and (*b*) *V_s_* ∪ *V_e_* induces a connected subgraph in *M*; (*iii*) computing a local phylogenetic tree *t* over *V_s_* ∪ *V_e_*; (*iv*) updating the global phylogenetic tree *T* by adding edges in subtrees of *t* that are induced by *V_s_* and (*v*) updating the MST and (*vi*) repeating steps (*ii*) to (*v*) until *T* is connected;

**Algorithm 1:**
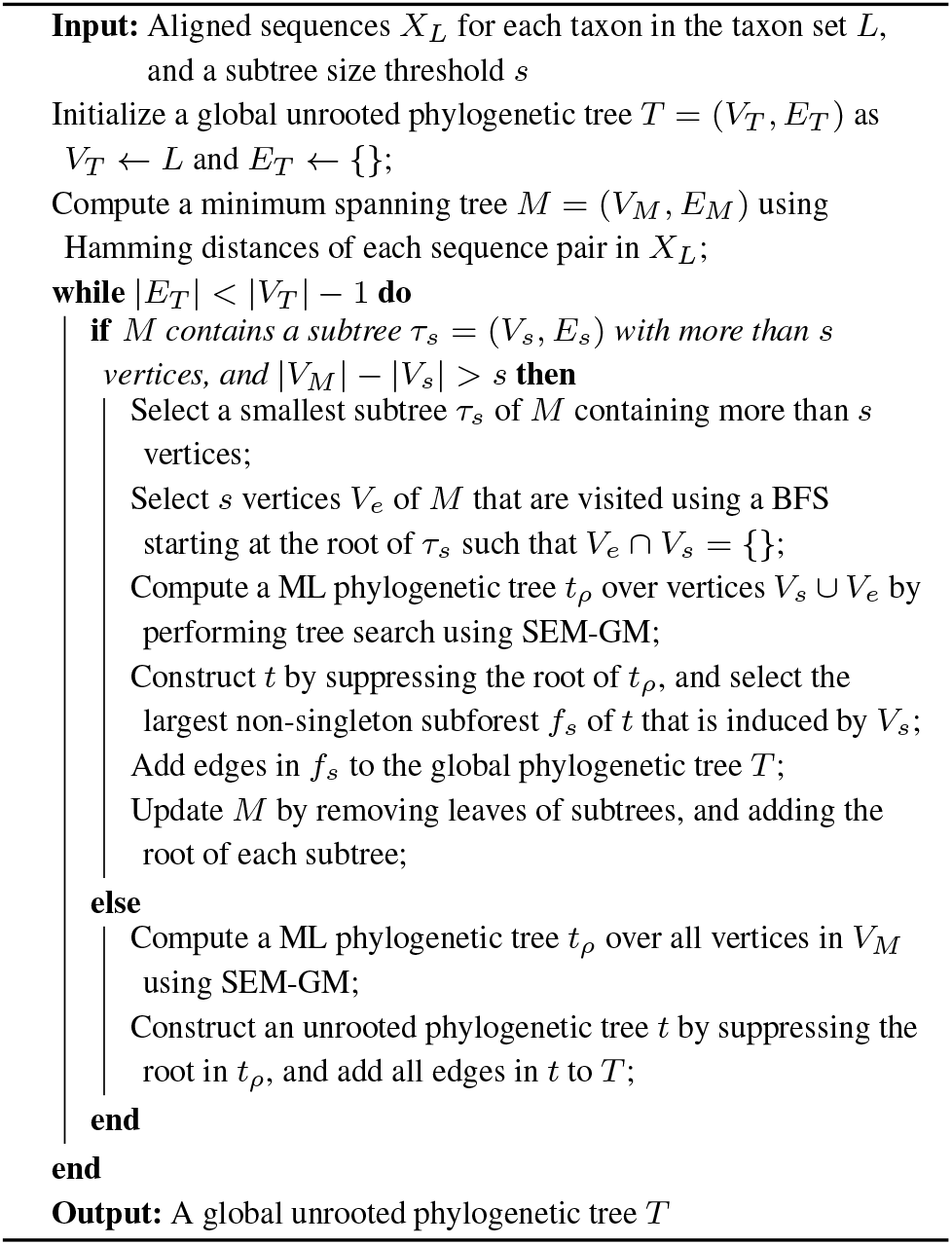
Constrained tree search using MST-backbone

Each step of MST-backbone is described in detail below. See Algorithm 1 for an overview of MST-backbone, and Fig. 2 for an illustration.

**Fig. 2.**
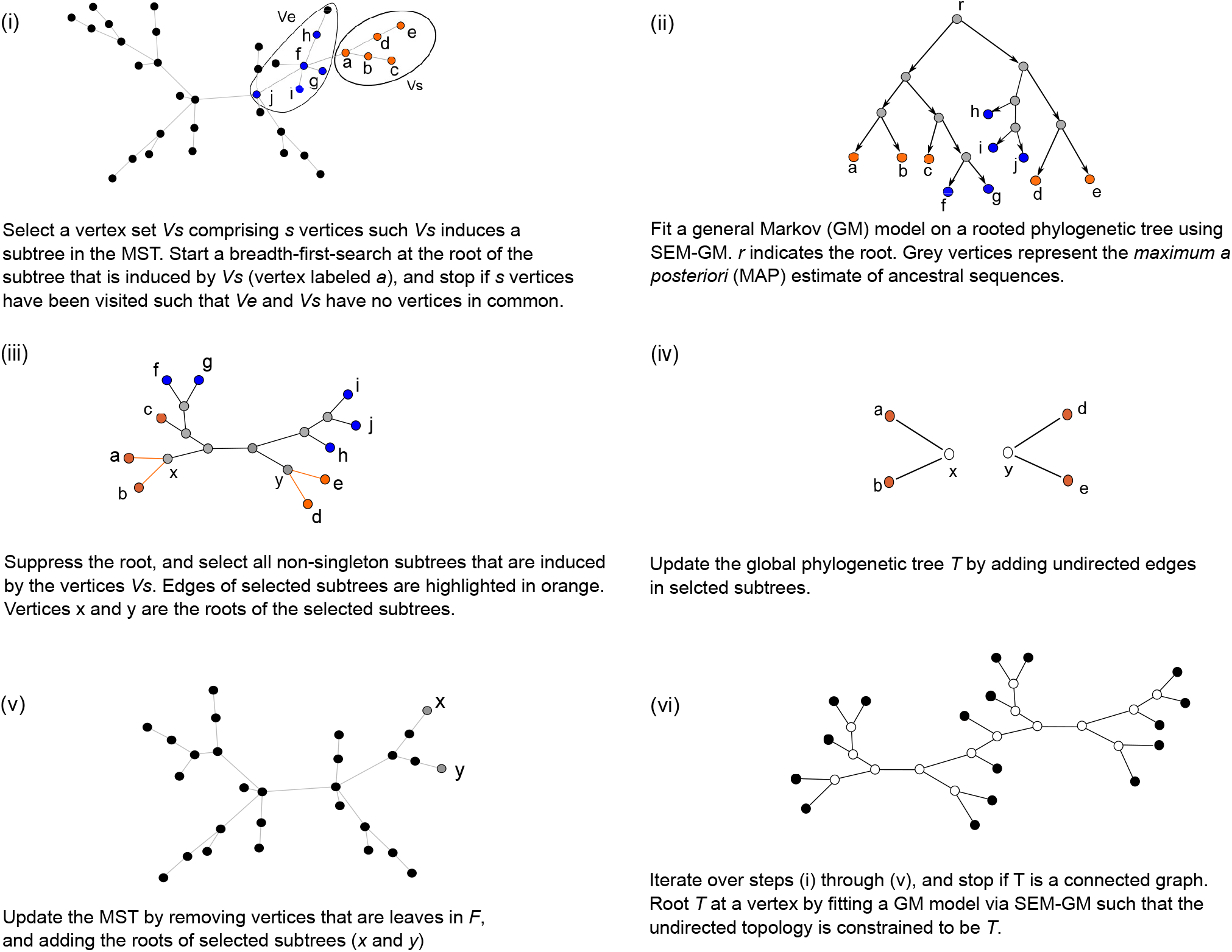
An illustration of MST-backbone

#### Initialization

A minimum spanning tree (MST) *M* is computed using the Hamming distance of each sequence pair in *X_L_*, where *L* represents the set of labeled vertices (species). We used Prim’s algorithm (Prim, 1957) for computing the initial MST. The global phylogenetic tree *T* = (*V_T_, E_T_*) is initialized by setting *V_T_* to *L*, and setting *E_T_* to the empty set.

#### Selecting subtrees of MST

Given a subtree size threshold *s*, select a smallest subtree *τ_s_* = (*V_s_, E_s_*) of *M* comprising more than *s* sequences. Subsequently, perform a breadth-first-search (BFS) on *M* starting at the root of *τ_s_*, and select *s* vertices *V_e_* such that *V_e_* and *V_s_* are mutually exclusive. We set the default value of the subtree size threshold to 10.

#### Computing local phylogenetic trees

An ML phylogenetic tree *t_ρ_* over *V_s_* ∪ *V_e_* is inferred using SEM-GM. A maximum a posteriori (MAP) sequence seq^MAP^(*h*) for each hidden vertex *h* is computed as 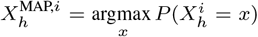 where 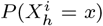 is the marginal probability for observing character *x* at site i for the sequence seq(*h*) that is represented by vertex *h*.

The location of the root as inferred in *t_ρ_* is not necessarily an optimal location of the root in the global phylogenetic tree. An unrooted phylogenetic tree *t* is constructed by suppressing the root in *t_ρ_*. Subsequently, the subforest 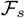 of *t* is selected such that (*i*) the leaves of each subtree in 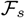 are in *V_s_*, and (*ii*) no component of 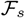 is a singleton vertex.

#### Updating the global phylogenetic tree

The global phylogenetic tree *T* = (*V_T_*, *E_T_*) is updated as follows. Non-leaf vertices of 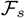 are added to *V_T_*. All edges of 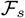 are added to *E_T_*. Edge length for each newly added edge {*u, v*} is computed as the normalized Hamming distance between sequence seq^MAP^(*u*) and seq^MAP^(*v*).

#### Updating the MST

Vertices that are leaves in 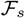 are removed from *V_M_*. The root of each subtree in 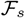 is added to *V_M_*. MAP estimates of sequences are used to compute the Hamming distances *d*(*r, v*) for each root *r* in 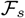, and each vertex *v* in *V_M_* ∩ {*V_s_* ∪ *V_e_*}. A new MST is computed using Prim’s algorithm using the newly computed distances.

### 2.4 Model selection

The General Markov model has the largest number of free parameters among DNA substitution models and may be prone to overtraining. We implemented a model selection framework called UNRESTselector for rooting trees inferred using MST-backbone(SEM-GM) under CT-HMMs that were parameterized in terms of one or more UNREST rate matrices. The number of distinct UNREST rate matrices was selected using BIC. Please refer to the supplementary section for more details.

### 2.5 Implementation details

We exponentiated UNREST matrices using the matrix exponential function as implemented in the Eigen 3 library (Guennebaud, Jacob, *et al.*, 2010). MSTs were computed using Prim’s algorithm as implemented in the Boost graph library (Siek *et al.*, 2000). Phylogenetic trees were simulated using the R package apTreeshape (Bortolussi *et al.*, 2006) and visualized using the R package ape (Paradis and Schliep, 2018). MST-backbone(SEM-GM) has been made available at https://github.com/prabhavk/mst-backbone-sem-gm under an open-source license. Custom python scripts were written in order to compute recall values, simulate sequences, clean alignments, and perform miscellaneous functions; the scripts have also been made available in the github repository mst-backbone-sem-gm.

## 3 Materials

### 3.1 Simulated data

We simulated non-stationary sequence evolution using a GM model in order to check for systematic error caused by using stationary homogeneous Markov models. Phylogenetic trees that were used for simulating sequence evolution were sampled from the uniform distribution over rooted trees using the R package apTreeshape v1.4.5 (Bortolussi *et al.*, 2006). A general Markov model was constructed for each phylogenetic tree as follows. The root probability *π_ρ_* was generated by sampling from the uniform distribution U(0, 1) and scaling such that the elements of *π_ρ_* summed to one. The transition matrices were generated as follows. The diagonal element of a transition matrix *P* was sampled from the uniform distribution U(*p*_min_, 1), where *p*_min_ was varied from 0.99 to 0.7. Smaller values of *p*_min_ result in greater change in GC content. Each non-diagonal element of *P* was sampled from the uniform distribution U(0, 1) and scaled such that each row of *P* summed to one. Sequence length was set to 1000 base pairs which is comparable to the number of columns in the empirical alignments that we analyzed which ranged from 128 bp to 2214 bp. We used the *χ*^2^ test to measure the extent to which base composition varies across taxa in simulated trees, because the *χ*^2^ test is widely used in practice. See Table 1 in the supplementary material for an overview of characteristics of the simulated data that was used in this study.

**Table 1.**
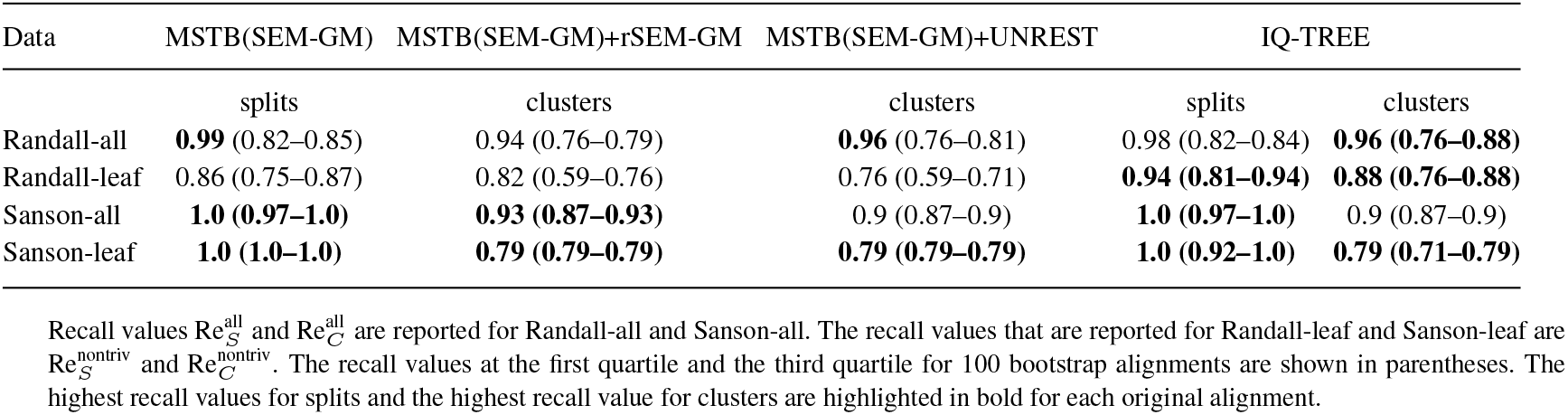
Recall values for experimental phylogenetic trees

| Data | MSTB(SEM-GM) | MSTB(SEM-GM)+rSEM-GM | MSTB(SEM-GM)+UNREST | IQ-TREE |  |
| --- | --- | --- | --- | --- | --- |
|  | splits | clusters | clusters | splits | clusters |
| Randall-all | **0.99** (0.82–0.85) | 0.94 (0.76–0.79) | **0.96** (0.76–0.81) | 0.98 (0.82–0.84) | **0.96 (0.76–0.88)** |
| Randall-leaf | 0.86 (0.75–0.87) | 0.82 (0.59–0.76) | 0.76 (0.59–0.71) | **0.94 (0.81–0.94)** | **0.88 (0.76–0.88)** |
| Sanson-all | **1.0 (0.97–1.0)** | **0.93 (0.87–0.93)** | 0.9 (0.87–0.9) | **1.0 (0.97–1.0)** | 0.9 (0.87–0.9) |
| Sanson-leaf | **1.0 (1.0–1.0)** | **0.79 (0.79–0.79)** | **0.79 (0.79–0.79)** | **1.0 (0.92–1.0)** | **0.79 (0.71–0.79)** |

Recall values $Re_S^{\text{all}}$ and $Re_C^{\text{all}}$ are reported for Randall-all and Sanson-all. The recall values that are reported for Randall-leaf and Sanson-leaf are $Re_S^{\text{nontriv}}$ and $Re_C^{\text{nontriv}}$ . The recall values at the first quartile and the third quartile for 100 bootstrap alignments are shown in parentheses. The highest recall values for splits and the highest recall value for clusters are highlighted in bold for each original alignment.

### 3.2 Empirical data

Two datasets were selected where GC content varied across taxa. The first data set comprises 13 beetle mitochondrial gene sequence alignments that were previously analyzed by Sheffield *et al.* (2009). The second data set comprises 1425 16S ribosomal RNA (16S rRNA) sequences from bacteria, archaea, and eukaryotes (Hug *et al.*, 2016). We subsampled the rRNA dataset to 100 sequences such that the proportion of bacterial, archaeal and eukaryotic sequences remained unchanged.

The true evolutionary history of genes is not known in general. We selected two experimental phylogeny data sets where gene sequences were evolved using the polymerase chain reaction according to a specified phylogenetic tree (Sanson *et al.* (2002), Randall *et al.* (2016)). The experimental phylogeny used by Randall et al. comprises 19 leaves and 330 ancestors, whereas the experimental phylogeny used by Sanson et al. comprises 16 leaves and 15 ancestors. Four alignments were constructed using the experimental phylogeny datasets: Randall-leaf and Sanson-leaf comprising leaf sequences, and Randall-all and Sanson-all comprising ancestor sequences and leaf sequences.

Additionally, we selected two virus data sets: HIV and Influenza A H3N2. The HIV dataset comprises 181 HIV *env* gene sequences that were sampled from 11 individuals that are part of a partially known transmission network (Vrancken *et al.*, 2014). The direction of transmission involving individuals A and B is not known. We downloaded all Influenza A H3N2 virus sequences from the GISAID data base (Shu and McCauley, 2017) with collection times ranging from 1968 to 2017. Subsequently, we discarded duplicate sequences and retained all earliest collected distinct sequences. Finally, we created a smaller data set by sampling at random at most five sequences per year of collection. The Influenza A H3N2 data set that was analyzed comprises 156 sequences.

We constructed multiple sequence alignments using MAFFT v7.3.3 (Katoh *et al.*, 2002; Katoh and Standley, 2013) and removed all alignment columns that contained gaps or ambiguous characters because. The size of the alignment constructed using MAFFT and the size of the trimmed alignment is shown in Table of the supplementary material. We quantified the extent to which empirical data violated the stationarity assumption using trimmed alignments (see Table 5 of supplementary material).

100 bootstrap alignments were constructed for each empirical alignment by sampling alignment columns with replacement. Bootstrap consensus trees were constructed by suppressing all vertices with bootstrap support less than 70%.

### 3.3 Measure of reconstruction accuracy

We measured reconstruction accuracy on simiulated data and the experimental phylogeny datasets by computing recall values for splits and clusters as defined below. 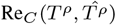 is the fraction of clusters in the model tree that are present in the estimated tree and is given by

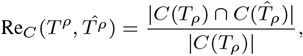

where *C*(*t_ρ_*) is the set of clusters in the rooted tree *t_ρ_*.

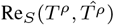 is the fraction of splits in the model tree that are present in the estimated tree and is give by

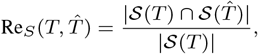

where 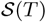 is the set of splits in the unrooted phylogenetic tree *T* that is constructed by suppressing the root of simulated tree *t_ρ_*.

A split is said to be a trivial split if the smallest side of the split contains one labeled vertex. A singleton cluster is said to be a trivial cluster.

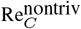 and 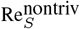 are recall values that are computed using nontrivial clusters and nontrivial splits, respectively. 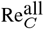 and 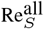 are recall values that are computed using all clusters, and all splits, respectively.

### 3.4 Statistical tests

The extent to which sequence alignments violated the stationarity assumption was quantified using a *χ*^2^ test with a p-value cutoff of 0.05. Statistical significance of difference in recall values was quantified using a t-test with a p-value cutoff of 0.05.

## 4 Results

### 4.1 Comparative analysis on empirical data

We were interested in investigating whether or not it is possible to infer realistically rooted trees using empirical data under the GM model. In addition to rooting trees under the GM model, we rooted trees via a CT-HMM that was selected using UNRESTselector. Additionally, we inferred rooted trees using IQ-TREE by selecting a stationary homogeneous CT-HMM using BIC. The models compared by IQ-TREE include the GTR model and submodels of the GTR, and Lie Markov models (a set of irreversible and time-reversible CT-HMM that are closed under matrix multiplication). The Lie Markov model 12.12 is the GM model (Woodhams *et al.*, 2015). However, model 12.12 as implemented in IQ-TREE is the UNREST model. In results presented below we refer to model 12.12 as UNREST. MST-backbone(SEM-GM) was run using a subtree size threshold of 10 because results on simulated data suggest that subtree size threshold does not significantly impact reconstruction accuracy.

#### 4.1.1 Results of model selection

The results of model selection are presented below (see Table 4 in the supplement for an overview). The Bayesian information criterion was used for selecting models. UNRESTselector chose one UNREST rate matrix for 17 out of 20 empirical alignments, and a CT-HMM with two UNREST rate matrices for 3 out of 20 empirical datasets. In the following text MST-backbone(SEM-GM)+UNREST refers to the method for rooting unrooted trees inferred using MST-backbone(SEM-GM) using the UNREST model. IQ-TREE selected irreversible models for 13/20 alignments and time-reversible models for 7/20 alignments (see Table in the supplement). We estimated rooted trees using IQ-TREE using the irreversible models that were selected by BIC. We inferred rooted trees using the UNREST+R2 model for the 7 datasets where IQ-TREE selected time-reversible models. R2 is the free-rate mixture model that allows substitution rates to vary across sites (Yang, 1995). See Table 4 of the supplementary material for an overview of the results of model selection.

#### 4.1.2 Experimental phylogenies

MST-backbone(SEM-GM) was able to recover the unrooted topology of experimental phylogenies with high recall values (0.99 or more for datasets comprising all sequences, and 0.86 or more for datasets comprising leaf sequences, see Table 1). MST-backbone(SEM-GM) outperformed IQ-TREE for the Randall-all dataset but performed worse on the Randall-leaf dataset. MST-backbone(SEM-GM) and IQ-TREE inferred the unrooted topology of the Sanson datasets with a recall of 1.0. The placement of the root wrt to the GM model was very accurate on datasets comprising all sequences with recall values of 0.93 and 0.94 for Sanson-all and Randall-all, respectively. Trees inferred using leaf sequences were rooted under the GM model with recall of 0.82 and 0.79 for Randall-leaf and Sanson-leaf, respectively. Trees rooted under the UNREST model scored higher recall values than trees rooted under the GM model for two datasets (Randall-all and Randall-leaf). Recall values for bootstrapped alignments are smaller than the recall values for the original alignment. For instance, 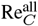 for IQ-TREE drops from 0.96 for the complete alignment for Randall-all to an inter-quartile range (IQR) of 0.76 to 0.88 for 100 bootstrapped alignments. The drop in recall for bootstrapped alignments is probably because the bootstrapped alignments provide less information because bootstrapped alignments almost always contain fewer distinct alignment columns compared to the original alignment.

#### 4.1.3 Beetle mitochondria

Phylogenetic trees for 13 beetle mitochondrial genes were estimated using MST-backbone(SEM-GM)+rSEM-GM and MST-backbone(SEM-GM)+UNRESTselector. Additionally, rooted trees were inferred using IQ-TREE. We validated the phylogenetic trees by counting the number of established taxonomic relationships that are compatible with the estimated trees. The established relationships among the beetles include (*i*) the monophyly of six species in the infraorder *Cucujiformia*, (*ii*) the monophyly of four species in the superfamily *Elateroidea*, and (*iii*) sister relationship between the suborders *Polyphaga* and *Archostemata* constituting, 12 species and 1 species, respectively.

MST-backbone(SEM-GM)+rSEM-GM, MST-backbone(SEM-GM)+ UNRESTselector, and IQ-TREE fared poorly in inferred realistic beetle mitochondrial phylogenies. None of the inferred gene trees are compatible with all established relationships. Four out of thirteen gene trees that were inferred using IQ-TREE are compatible with one out of three established relationships each. Only one gene tree inferred by MST-backbone(SEM-GM)+rSEM-GM and only one gene tree inferred by MST-backbone(SEM-GM)+UNRESTselector was compatible with one out of three established relationships (see Table 2).

**Table 2.**
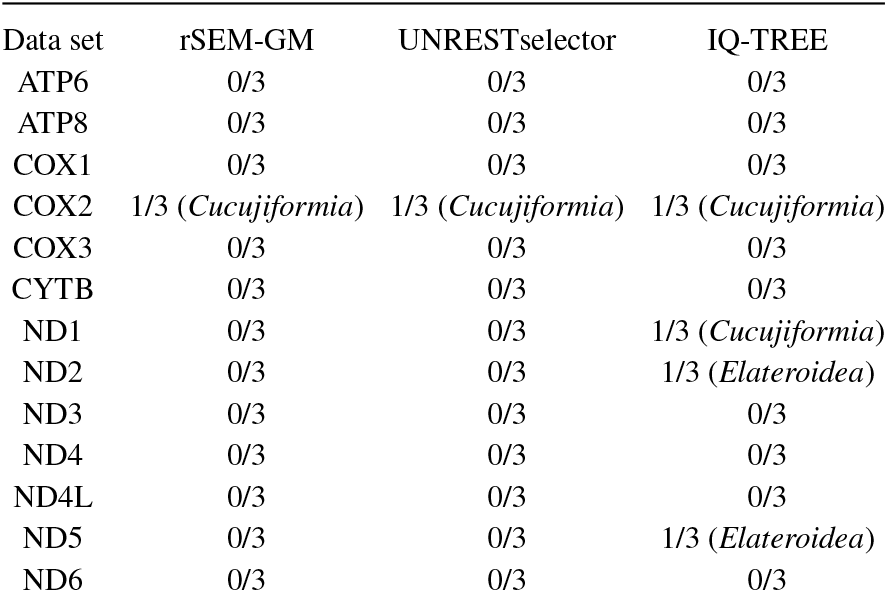
Number of established evolutionary relationships that are compatible with inferred trees

| Data set | rSEM-GM | UNRESTselector | IQ-TREE |
| --- | --- | --- | --- |
| ATP6 | 0/3 | 0/3 | 0/3 |
| ATP8 | 0/3 | 0/3 | 0/3 |
| COX1 | 0/3 | 0/3 | 0/3 |
| COX2 | 1/3 (*Cucujiformia*) | 1/3 (*Cucujiformia*) | 1/3 (*Cucujiformia*) |
| COX3 | 0/3 | 0/3 | 0/3 |
| CYTB | 0/3 | 0/3 | 0/3 |
| ND1 | 0/3 | 0/3 | 1/3 (*Cucujiformia*) |
| ND2 | 0/3 | 0/3 | 1/3 (*Elateroidea*) |
| ND3 | 0/3 | 0/3 | 0/3 |
| ND4 | 0/3 | 0/3 | 0/3 |
| ND4L | 0/3 | 0/3 | 0/3 |
| ND5 | 0/3 | 0/3 | 1/3 (*Elateroidea*) |
| ND6 | 0/3 | 0/3 | 0/3 |

#### 4.1.4 Ribosomal RNA

The unrooted tree inferred by MST-backbone(SEM-GM) and the unrooted version of the phylogenetic tree inferred by IQ-TREE are compatible with the three-domain classification proposed by Woese *et al.* (1990). Bacteria, archaea and eurkaryotes formed unrooted subtrees with bootstrap support greater than 95% for MST-backbone(SEM-GM) and the unrooted version of trees inferred by IQ-TREE (see Figure 4 A in the supplementary material). The root is located among bacteria for trees rooted under the GM model and the UNREST model that was selected by UNRESTselector. The trees that were inferred using IQ-TREE under the UNREST+R7 model were rooted among archaea (see Figure 4 B in the supplementary material). A realistic rooting would be one were the least common ancestor of bacteria would be a direct descendant of the root, assuming that ribosomes in extant cells have evolved from ribosomes present in the last universal common ancestor.

### 4.2 Influenza A H3N2

Under the assumption of a strict molecular clock, the number of character changes that accumulate in a sequence are proportional to the collection time of the sequence. The Influenza A H3N2 virus exhibits a strict clock-like evolution (Gojobori *et al.*, 1990). We validated inferred phylogenetic trees by measuring the Pearson’s correlation coefficient of collection times with weighted root-to-leaf path lengths in inferred phylogenetic trees.

The phylogenetic tree that was inferred under the GM model has a Pearson’s correlation coefficient of −0.92 indicating that the location of the root is not realistic (see Figure 3 A). Collection times are highly correlated with root-to-leaf path lengths in the phylogenetic tree that was inferred with MST-backbone(SEM-GM) and rooted under the UNREST model (Pearson’s *ρ* of 0.989; see Figure 3 B). The collection times are highly correlated with root-to-leaf path lengths with Pearson’s *ρ* of 0.996 in the phylogenetic tree inferred using IQ-TREE under the UNREST + R2 model (see Figure 3 C). Correlation scores for bootstrap consensus trees are high (*ρ* of 0.989 for MST-backbonse(SEM-GM)+UNREST and *ρ* of 0.996 for IQ-TREE using UNREST +R2, see Figure 1 in the supplement material)

**Fig. 3.**
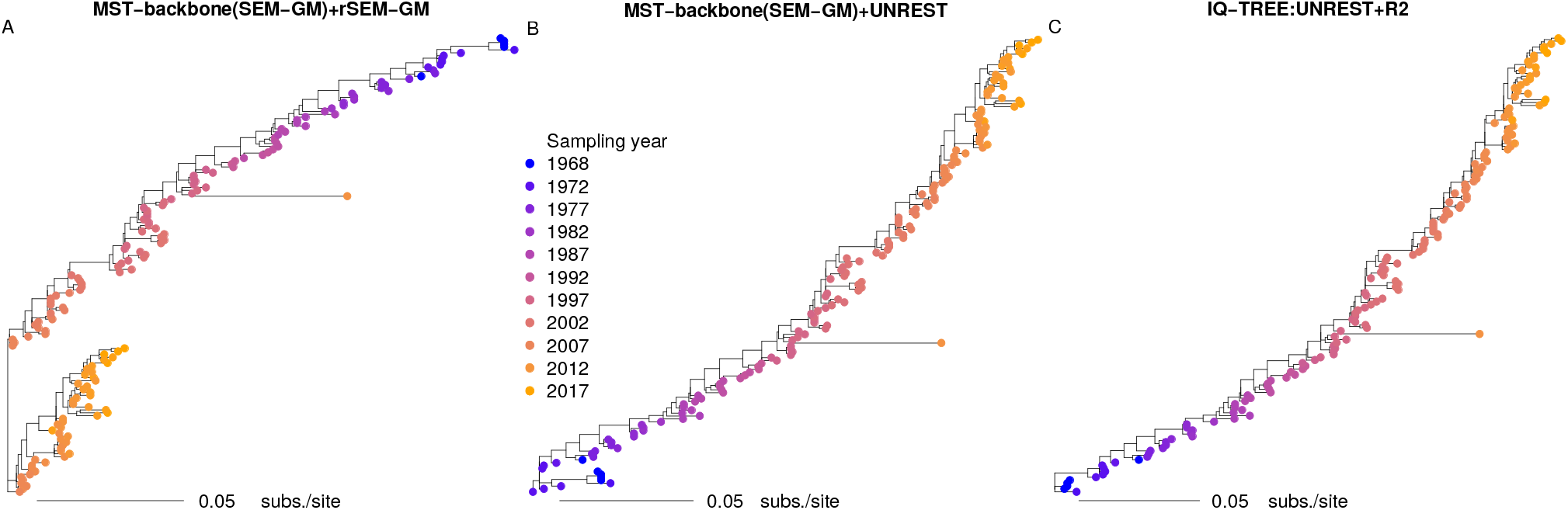
H3N2 phylogenetic trees that were inferred using MST-backbone (SEM-GM), and subsequently rooted under the GM model (panel A) and the UNREST model. An H3N2 phylogenetic tree that was inferred using IQ-TREE under the UNREST+R2 model is shown in panel C

### 4.3 HIV transmission network

A rooted pathogen phylogenetic tree is said to be compatible with a transmission event if pathogens from the recipient have descended from pathogens of the transmitter. The phylogenetic tree for HIV that was inferred by MST-backbone(SEM-GM)+rSEM-GM is not rooted realistically because it is rooted at a sequence from individual C which is not compatible with the transmission from B to C (see Figure 4 A). We rooted the tree that was inferred using MST-backbone(SEM-GM) under the UNREST model, and found that the rooted tree is compatible with nine out of ten transmission events (see Figure 4 B). The transmission event *B* → *I* is not compatible with the phylogenetic tree. We performed model selection using IQ-TREE and found that IQ-TREE selected the time-reversible model TVM+F+R4. The rooted phylogenetic tree inferred using IQ-TREE with the UNREST+R2 is compatible with all transmission events (see Figure 4 C).

**Fig. 4.**
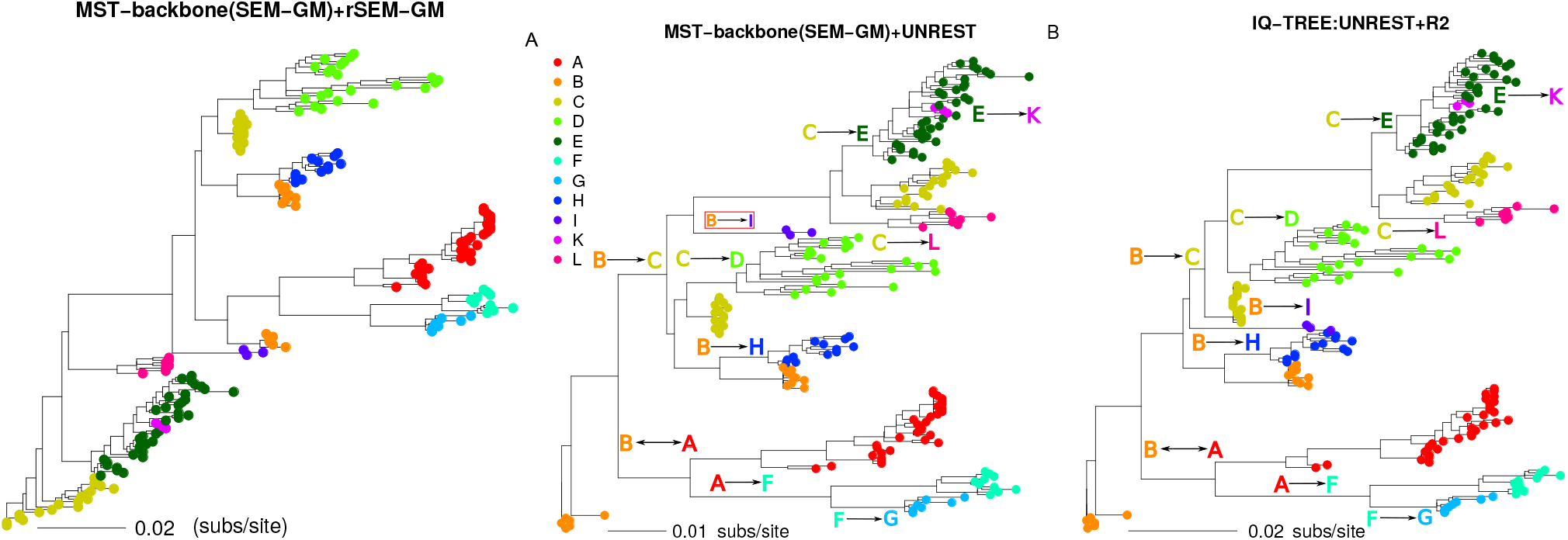
HIV phylogenetic trees that were inferred using MST-backbone (SEM-GM), and subsequently rooted under the GM model (panel A) and the UNREST model. An H3N2 phylogenetic tree that was inferred using IQ-TREE under the UNREST+R2 model is shown in panel C. Leaves are coloured wrt infected host.

Bootstrap consensus trees for MST-backbone(SEM-GM)+UNREST and IQ-TREE are poorly resolved (see Figure 2 in the supplementary material). Only 27% and 24% of vertices in bootstrap consensus trees have bootstrap support greater than 70%, for MST-backbone(SEM-GM)+UNREST and IQ-TREE, respectively.

### 4.4 Comparative analysis on simulated data

We compared MST-backbone(SEM-GM) and MST-backbone(SEM-GM)+rSEM-GM with IQ-TREE v2.0 (Minh *et al.*, 2020), RAxML-NG v0.8.1 (Kozlov *et al.*, 2019) and FastTree v2.1.10 (Price *et al.*, 2010), which are among the most widely used software for inferring ML phylogenetic trees. FastTree and RAxML-NG perform phylogeny inference using time-reversible models, whereas the DNA substitution models implemented in IQ-TREE include time-reversible and non-reversible Markov models. FastTree and RAxML-NG were executed under the GTR model with the number of rate categories set to one. RAxML-NG requires the user to specify the number of starting trees and the type of starting tree (random or parsimony). We ran RAxML-NG with one parsimony starting tree. IQ-TREE was used to infer rooted trees using the UNREST model.

We report two recall values for simulated data: 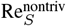 and 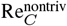, which denote the fraction of non-trivial splits and non-trivial clusters in simulated trees that are present in estimated trees, respectively (see Subsection 3.3).

We measured branch length for each setting of *p*_min_ in order to facilitate comparison with simulation experiments that are performed using continuous-time Markov models. Branch length was computed as the number of observed substitutions per site. Average branch lengths ranged from 0.0025 subs/site (*p*_min_ = 0.995) to 0.15 subs/site (*p*_min_ = 0.7), see Table 4. Average branch lengths for empirical alignments ranged from 0.001 subs/site (Sanson-all) to 0.12 subs/site (*ND2* gene) (see Table in supplementary material).

MST-backbone(SEM-GM) scored significantly higher 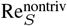 values (*p* < 0.05) compared to competing methods for *p*_min_ values smaller than 0.9 (see Table 3). All methods performed similarly for *p*_min_ values ranging from 0.9 to 0.995; i.e., the pairwise difference in mean 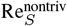 values was not statistically significant for *p*-value cut-off of 0.05. In general, all methods score median 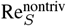 values of 0.98 or higher for *p*_min_ ranging from 0.9 to 0.99.

**Table 3.** 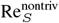 values for inferring unrooted topology of simulated trees

| $p_{\min}$ | MST-backbone(SEM-GM) | FastTree | RAxML-NG | IQ-TREE |
| --- | --- | --- | --- | --- |
| 0.7 | **0.88 (0.014)** | 0.81 (0.035) | 0.75 (0.04) | 0.82 (0.048) |
| 0.8 | **0.95 (0.009)** | 0.87 (0.044) | 0.9 (0.03) | 0.92 (0.06) |
| 0.9 | 0.99 (0.004) | **1.0 (0.0)** | **1.0 (0.0)** | **1.0 (0.001)** |
| 0.95 | **1.0 (0.001)** | **1.0 (0.0)** | **1 (0.0)** | **1.0 (0.001)** |
| 0.99 | 0.98 (0.0045) | **0.99 (0.005)** | 0.98 (0.004) | 0.98 (0.005) |
| 0.995 | **0.92 (0.013)** | **0.92 (0.013)** | **0.92 (0.013)** | **0.92 (0.012)** |

The significantly highest $\text{Re}_S^{\text{nontriv}}$ values in each row are highlighted in bold. Values shown within parentheses are interquartile ranges.

**Table 4.** 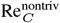 values for inferring rooted topology of simulated trees

| $p_{\min}$ | average branch length | % seqs with $p < 0.05$ for $\chi^2$ test | $\overline{\Delta_f}$ | MST-backbone(SEM-GM)+rSEM-GM | IQ-TREE |
| --- | --- | --- | --- | --- | --- |
| 0.7 | 0.15 (0.064) | 89.8 (0.8) | 0.1 | **0.83 (0.033)** | 0.8 (0.049) |
| 0.8 | 0.1 (0.043) | 83.2 (2.5) | 0.068 | **0.9 (0.045)** | **0.9 (0.054)** |
| 0.9 | 0.05 (0.022) | 68.0 (8.3) | 0.035 | 0.93 (0.038) | **0.96 (0.037)** |
| 0.95 | 0.025 (0.012) | 53.5 (19.9) | 0.02 | 0.94 (0.037) | **0.96 (0.033)** |
| 0.99 | 0.005 (0.004) | 19.9 (34.8) | 0.006 | 0.93 (0.035) | **0.94 (0.032)** |
| 0.995 | 0.0025 (0.0014) | 2.5 (19.1) | 0.0035 | 0.87 (0.044) | **0.88 (0.029)** |

$\text{Re}_C^{\text{nontriv}}$ values are listed in the columns indexed "MST-backbone(SEM-GM)+rSEM-GM" and "IQ-TREE". The significantly highest $\text{Re}_C^{\text{nontriv}}$ values in each row are highlighted in bold. Values shown within parentheses are interquartile ranges.

Rooted trees inferred using MST-backbone(SEM-GM)+rSEM-GM scored higher recall values (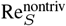 of 0.88) than IQ-TREE for *p*_min_ value of 0.7. Rooted trees inferred using IQ-TREE scored higher Re_*C*_ values for *p*_min_ ranging from 0.9 to 0.995. Recall values are low for large branches (0.1 subs/site or higher) because of increased odds of multiple substitutions at the same site. Recall values are low on the other end of the spectrum of edge lengths (0.0025 subs/site or smaller) because there are branches across which no substitutions have occurred.

We compared how recall values vary with the number of sequences that exhibit statistically significant deviation in base composition from the average base composition. On average, at least 68% of sequences scored significantly on the *χ*^2^ test for *p*_min_ values of 0.9 or smaller. At least 53.5% of sequences and 19.9% of sequences, with an inter-quartile range of 19.9% and 35.8%, scored significantly for *p*_min_ setting of 0.95 and 0.99. The number of sequences that score significantly on the *χ*^2^ test does not seem to be a good indicator of reconstruction accuracy. *p*_min_ or average branch length seem to be a better indicator of reconstruction accuracy.

MST-backbone(SEM-GM)+rSEM-GM scored significantly higher 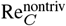 values compared to IQ-TREE for *p*_min_ of 0.7. IQ-TREE had highest 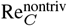 for *p*_min_ values ranging from 0.9 to 0.995 (see Table 4), suggesting that the UNREST model may be more accurate than the GM model at placing the root for trees with edges shorter than 0.05 subs/site.

We measured the effect of subtree size threshold on the performance of MST-backbone(SEM-GM) by varying subtree threshold size from 10 to 40. 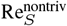 values did not vary significantly for varying threshold size (see Table 2 in supplementary material).

#### 4.4.1 Scalability

In order to perform a comparative analysis of scalability we varied the number of taxa ranging from 1000 to 5000 and set *p*_min_ to 0.99, and measured elapsed CPU times. We chose a setting of 0.99 for *p*_min_ because GM models generated by setting *p*_min_ to 0.99 correspond to CT-HMM with average branch length of 0.005 subs/site, which lies within the range (0.001 subs/site to 0.12 subs/site) of the average branch length of empirical phylogenetic trees that were inferred using MST-backbone(SEM-GM). We split the elapsed CPU time for MST-backbone(SEM-GM)+rSEM-GM into time taken by MST-backbone(SEM-GM) to infer a global unrooted tree, and the time taken by rSEM-GM to root a global unrooted tree using the GM model. Running SEM-GM without using MST-backbone turned out to be extremely computationally expensive and was only performed for trees with 1000 taxa. MST-backbone(SEM-GM) and FastTree appear to scale linearly whereas rSEM-GM, IQ-TREE and RAxML-NG appear to scale quadratically (see Figure 5).

**Fig. 5.**
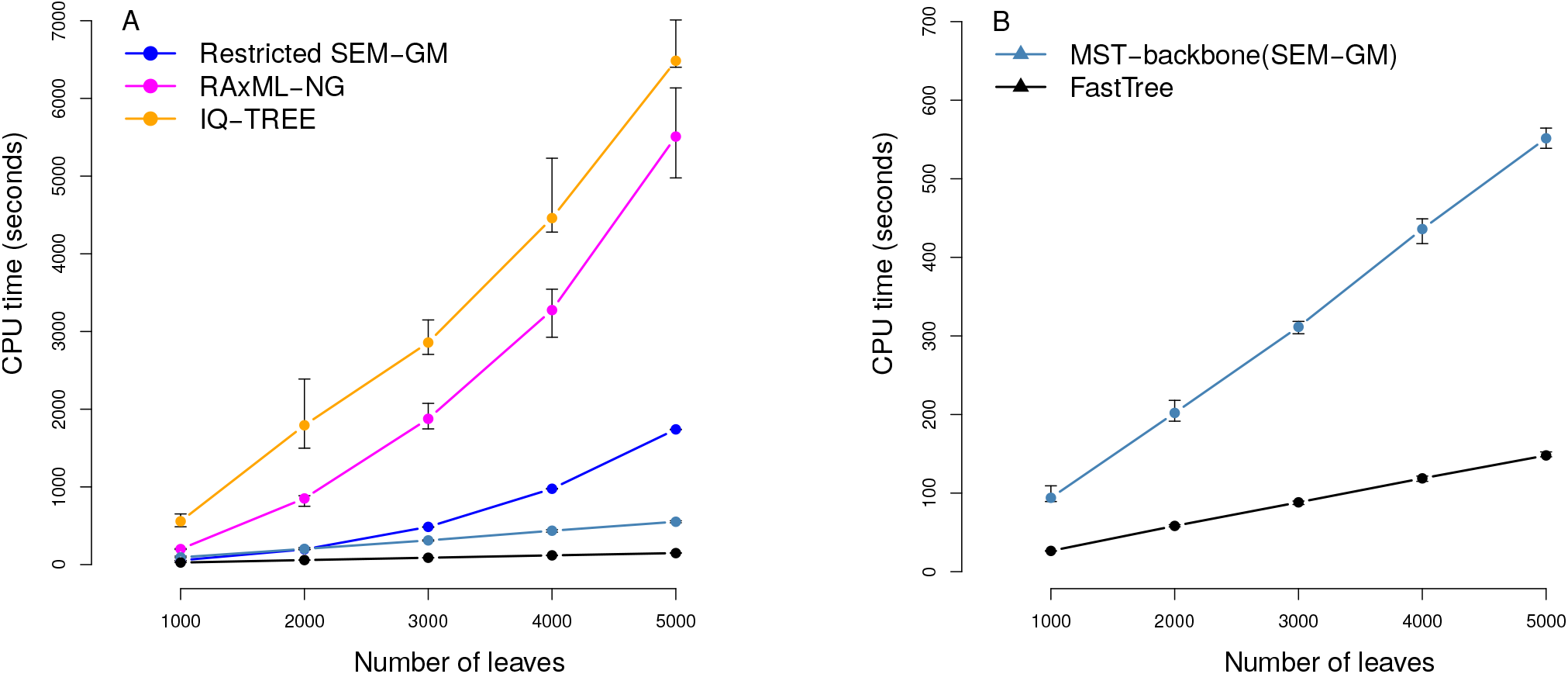
Elapsed CPU times for MST-backbone(SEM-GM), MST-backbone(SEM-GM)+rSEM-GM, RAxML-NG, IQ-TREE and FastTree. Simulated sequences were generated by setting *p*_min_ to 0.99. Error bars represent IQR computed using 20 replicates.

RAxML-NG and IQ-TREE optimize the parameters of CT-HMM by alternatively optimizing a rate matrix for fixed tree topology and branch lengths, and optimizing branch lengths subsequent to each tree rearrangement operation for a fixed rate matrix, until the log-likelihood score converges. Rate matrices are optimized using numerical procedures that require repeated tree traversals, with each traversal costing *O*(*nkA*^2^) floating-point operations, where *n* is the number of taxa, *k* is the sequence length, and *A* is the number of states (four for DNA). RAxML-NG and IQ-TREE reuse conditional likelihood vectors subsequent to tree rearrangement operations such as nearest neighbor interchange (NNI) and subtree prune and regraft (SPR) in order to optimize branch lengths. Each expectation step of rSEM-GM requires *O*(*nkA*^2^) floating-point operations, whereas each maximization step requires *O*(*nA*^2^) floating-point operations. The quadratic scaling of RAxML-NG, IQ-TREE and rSEM-GM is probably because the number of tree traversals required for the log-likelihood score to converge grows linearly in number of taxa.

The CPU-time consumed by IQ-TREE and RAxML-NG is at least 3.16 times the CPU-time consumed by rSEM-GM because IQ-TREE and RAxML-NG perform extensive tree-search combined with parameter optimization whereas rSEM-GM roots an unrooted tree under the GM model. SEM-GM consumed around 24 hours of CPU time for inferring trees with 1000 leaves if tree search was not constrained using MST-backbone.

MST-backbone(SEM-GM) scales linearly because the most CPU intensive task is to infer local trees using SEM-GM, and the number of local trees grows roughly linearly with number of taxa.

FastTree uses two optimization scores, minimum evolution (ME) and maximum likelihood. ME trees are phylogenetic trees with the shortest tree length (sum of edge lengths), where edge lengths are obtained by regressing weighted path lengths on pairwise leaf-to-leaf distances (Desper and Gascuel, 2002). FastTree uses a large number of heuristics that constrain search through tree-space and reduce the cost of parameter optimization. FastTree computes a starting tree using NJ and subsequently searches for ME trees by using NNI moves and linear SPR moves, i.e., SPR moves that can be performed in *O*(*n*) steps. Subsequent to the ME phase, FastTree searches for ML trees using a limited number of NNI moves. FastTree considerably saves on time spent in parameter optimization by optimizing a GTR rate matrix once. Additionally FastTree uses conditional likelihood vectors to performs branch length optimization. FastTree makes use of a faster-to-compute score that approximates the likelihood score during the search for ML trees. It may be that FastTree scales linearly on simulated data because “linear SPR” based ME search is sufficient to obtain a good estimate of tree topology and branch lengths. FastTree is around 3.6 times faster than MST-backbone(SEM-GM).

## 5 Discussion

Widely used software for model-based phylogeny inference makes simplifying assumptions about gene evolution such as stationarity, homogeneity and time-reversibility. The wide-spread use of stationary models is thought to have led to systematic error in inferred phylogenies (Sheffield *et al.*, 2009). The general Markov model (GM) is an alternative to the continuous-time Markov models that are widely used for phylogeny inference. Pachter and Sturmfels (2005) note that the GM model can optimized using the expectation-maximization approach. In the current study we performed tree search under the GM model by applying the structural EM algorithm (SEM; Friedman *et al.* (2002)) to the GM model. Additionally, we implemented a minimum spanning tree framework called MST-backbone for constraining search through tree-space because structural EM is computationally expensive.

Results for five out of six empirical data sets suggest that the unrooted topology can be recovered reliably using MST-backbone(SEM-GM). Only a small fraction of mitochondrial gene trees inferred by MST-backbone(SEM-GM)+rSEM-GM, MST-backbone(SEM-GM)+ UNRESTselector and IQ-TREE is compatible with established taxonomic relationships. The mitochondrial gene sequences studied by us were previously analyzed by Sheffield *et al.* (2009). Sheffield *et al.* (2009) inferred trees using a concatenated alignment of gene sequences, and found that consensus trees inferred using p4 and PHASE were compatible with all established relationships, whereas the trees inferred using nhPhyML were compatible with established relationships only if the starting tree was the consensus tree that was inferred by p4.

The location of the root as inferred by rSEM-GM is accurate for two experimental phylogeny datasets and the simulated datasets. Rooting the trees inferred by MST-backbone(SEM-GM) under the UNREST model resulted in realistically rooted trees for two experimental phylogeny datasets and two virus datasets but not for beetle mitochondria dataset or the 16S ribosomal RNA dataset. Similar results were obtained using IQ-TREE which was run using the UNREST model and other time-irreversible Lie Markov models.

A comparative analysis on simulated data suggests that MST-backbone(SEM-GM) and FastTree scale linearly with number of leaves whereas rSEM-GM, IQ-TREE and RAxML-NG appear to scale quadratically with number of leaves.

In conclusion, we have presented SEM-GM a new method for performing tree search under the GM model, and we presented MST-backbone, an easy-to-implement framework for constraining search through tree space. Extensive validation on empirical data suggests that it may be possible to quickly infer unrooted trees using MST-backbone(SEM-GM) and it may be possible to infer realistically rooted trees using the UNREST model. It does not seem necessary to use the GTR model for computational ease.

## Acknowledgements

I thank Thomas Lengauer for reading this manuscript and for helpful discussion. I thank Elizabeth Allman and John Rhodes for suggesting that I apply the MST-backbone framework to the general Markov model. I thank Martin Vingron for helpful discussion

