## Supplementary material for "Phylogeny inference under the general Markov model using MST-backbone"

### Supplementary material for Phylogeny inference under the general Markov model using a minimum spanning tree backbone

Prabhav Kalaghatgi

The notation used in the supplement follows the terminology introduced in the main paper.

#### 1 Transforming a non-bifurcating generally labeled phylogenetic tree to a bifurcating leaf-labeled phylogenetic tree

Let $T_\rho$ be a rooted phylogenetic tree that does not have a bifurcating topology, and let $M$ be the GM model on $T_\rho$ that is computed by a maximization step of SEM-GM. We transform $T_\rho$ into a bifurcating tree $T_\rho^{\text{bi}}$, and compute a GM model on $T_\rho^{\text{bi}}$ such that the log-likelihood score remains unchanged. The transformation operations are described in Algorithm 1. We use conditional likelihood vectors (Felsenstein, 1981) in order to show that the likelihood score for any site $i$ remains unchanged subsequent to the operations applied for each case considered by Algorithm 1.

We assume that alignment columns $X^i$ are independently drawn from a common distribution (iid). The likelihood for site $i$ is defined as

$$L^i = \sum_x \pi_\rho(x) L^i_\rho(x)$$

where the conditional likelihood vector $L^i_v$ for each hidden vertex is given by

$$L^i_u(x) = \prod_{c \in \mathcal{C}(v)} \left( \sum_y P_{(u,c)}(y|x) L^i_c(y) \right),$$

where $P_{(u,c)}$ is the conditional probability $P(X_u|X_c)$.

$L^i_u(x)$ is the marginal probability

$$L^i_u = \sum_{X^i_h : h \in H_{\tau_u} \setminus \{u\}} P(\{X^i_v : v \in V_{\tau_u}\}, M_{\tau_u}),$$

where $\tau_u$ is the subtree of $T_\rho$ that is rooted at $u$. $M_{\tau_u}$ represents the set of transition matrices $\{P_e : e \in \tau_u\}$

The conditional likelihood vector for a labeled leaf vertex $l$ is defined as

$$L^i_l(x) = \delta(x, X^i_l),$$

where $\delta(x, y)$ is the Kronecker-delta function that equals one if $x$ equals $y$, and zero otherwise.

The conditional likelihood vector $L^i_u$ for a labeled non-leaf vertex $u$ is defined as

$$L^i_u(x) = \delta(x, X^i_u) \prod_{c \in C(u)} \left( \sum_y P_{(u,c)}(y|x) L^i_c(y) \right),$$

where $C(u)$ are the children of $u$.

The proof of correctness of Algorithm 1 is provided below.

**Lemma 1.** *The output of Algorithm 1 is a GM model on a bifurcating leaf-labeled phylogenetic tree such that log-likelihood remains unchanged.*

*Proof.* Note that the removal of edges incident to hidden vertices in cases $(i)$ through $(iii)$ results in the construction of the singleton hidden vertices that are used in cases $(iv)$ through $(vi)$.

**Case** $(i)$**:** $T_\rho$ contains a hidden leaf $h$.

Let $\mathcal{C}(v)$ be the set of children of the parent $v$ of $h$. The conditional likelihood vector $L_v^i$ is computed as follows.

$$L_v^i(x) = \left(\sum_y P_{(v,h)}(y|x)L_h^i(y)\right) \prod_{c\in\mathcal{C}(v)\setminus\{h\}} \left(\sum_z P_{(v,c)}(z|x)L_c^i(z)\right) \tag{1}$$

$$= \left(\sum_y P_{(v,h)}(y|x)\right) \prod_{c\in\mathcal{C}(v)\setminus\{h\}} \left(\sum_z P_{(v,c)}(z|x)L_c^i(z)\right) \quad (L_h^i(y) \text{ equals one for all } y \text{ because } h \text{ is not observed}) \tag{2}$$

$$= \prod_{c\in\mathcal{C}(v)\setminus\{h\}} \left(\sum_z P_{(v,c)}(z|x)L_c^i(z)\right) \quad (\text{Each row of } P_{(v,h)} \text{ sums to one}) \tag{3}$$

**Case** $(ii)$**:** $T_\rho$ contain a hidden vertex $h$ with in-degree one and out-degree one.

Let $u$ and $v$ be the parent and child, respectively, of $h$. The conditional likelihood vector $L_u^i$ is computed as follows

$$L_u^i(x) = \left(\sum_y P_{(u,h)}(y|x)L_h^i(y)\right) \prod_{c\in\mathcal{C}(u)\setminus\{h\}} \left(\sum_z P_{(u,c)}(z|x)L_c^i(z)\right) \tag{4}$$

$$= \left(\sum_y P_{(u,h)}(y|x)\sum_w P_{(h,v)}(w|y)L_v^i(w)\right) \prod_{c\in\mathcal{C}(u)\setminus\{h\}} \left(\sum_z P_{(u,c)}(z|x)L_c^i(z)\right) \tag{5}$$

$$= \left(\sum_w \left(\sum_y P_{(h,v)}(w|y)P_{(u,h)}(y|x)\right) L_v^i(w)\right) \prod_{c\in\mathcal{C}(u)\setminus\{h\}} \left(\sum_z P_{(u,c)}(z|x)L_c^i(z)\right) \tag{6}$$

$$= \left(\sum_w P_{(u,v)}(w|x)L_v^i(w)\right) \prod_{c\in\mathcal{C}(u)\setminus\{h\}} \left(\sum_z P_{(u,c)}(z|x)L_c^i(z)\right) \quad (\text{where } P_{(u,v)} = P_{(u,h)}P_{(h,v)}) \tag{7}$$

**Case** $(iii)$**:** The root is a hidden vertex with out-degree one.

Let $\pi_\rho^{\text{cur}}$ denote the current root probability distribution. The likelihood $L^i$ for site $i$ is computed as

$$L^i = \left(\sum_x \pi_\rho^{\text{cur}}(x)L_\rho^i(x)\right) \tag{8}$$

$$= \left(\sum_x \pi_\rho^{\text{cur}}(x)\sum_y P_{(\rho,v)}(y|x)L_v^i(y)\right) \tag{9}$$

$$= \left(\sum_y \sum_x \pi_\rho^{\text{cur}}(x)P_{(\rho,v)}(y|x)L_v^i(y)\right) \tag{10}$$

$$= \left(\sum_y \pi_\rho^{\text{new}}(y)L_v^i(y)\right) \quad (\text{where } \pi_\rho^{\text{new}}(y) = \sum_x \pi_\rho^{\text{cur}}(x)P_{(\rho,v)}(y|x)) \tag{11}$$

**Case** $(iv)$**:** $T_\rho$ contains a non-leaf labeled vertex $l$.

$$L^i_l(x) = \delta(x, X^i_l) \prod_{c \in \mathcal{C}(l)} \left( \sum_z P_{(l,c)}(z|x) L^i_c(z) \right) \tag{12}$$

$$= \delta(x, X^i_l) \prod_{c \in \mathcal{C}(h)\setminus\{l\}} \left( \sum_z P_{(h,c)}(z|x) L^i_c(z) \right), \tag{13}$$

$$\text{where } \delta() \text{ is the Kroenecker delta function} \tag{14}$$

$$\tag{15}$$

Consider the conditional likelihood vector $L^i_h$ of the hidden vertex $h$.

$$L^i_h(x) = \prod_{c \in \mathcal{C}(h)} \left( \sum_z P_{(h,c)}(z|x) L^i_c(z) \right) \tag{16}$$

$$= \left( \sum_y P_{(h,l)}(y|x) L^i_l(y) \right) \prod_{c \in \mathcal{C}(h)\setminus\{l\}} \left( \sum_z P_{(h,c)}(z|x) L^i_c(z) \right) \tag{17}$$

$$= \left( \sum_y P_{(h,l)}(y|x) \delta(y, X^i_l) \right) \prod_{c \in \mathcal{C}(h)\setminus\{l\}} \left( \sum_z P_{(h,c)}(z|x) L^i_c(z) \right) \tag{18}$$

$$\text{(because } L^i_l(y) = \delta(y, X^i_l) \text{ for labeled leaves)} \tag{19}$$

$$= P_{(h,l)}(x|x) \delta(x, X^i_l) \prod_{c \in \mathcal{C}(h)\setminus\{l\}} \left( \sum_z P_{(h,c)}(z|x) L^i_c(z) \right) \tag{20}$$

$$\text{(because } P_{(h,l)} \text{ is the identity matrix )} \tag{21}$$

$$= \delta(x, X^i_l) \prod_{c \in \mathcal{C}(h)\setminus\{l\}} \left( \sum_z P_{(h,c)}(z|x) L^i_c(z) \right) \tag{22}$$

The conditional likelihood vectors $L^i_h$ and $L^i_l$ are identical (see equation (13) and equation (22)).

**Case** $(v)$**:** $T_\rho$ contains a hidden vertex $h_1$ with out-degree greater than two. Let $C_1$ be the set of all children of $h_1$ prior to the operations performed in case $(v)$. Let $C_{uv}$ be $C_1\setminus\{u, v\}$.

The conditional likelihood vector $L^i_u$ prior to transformation operations is given by

$$L^i_{h_1}(x) = \left( \sum_y P_{(h_1,u)}(y|x) L^i_u(y) \right) \left( \sum_y P_{(h_1,v)}(y|x) L^i_v(y) \right) \prod_{d \in D_{uv}} \left( \sum_z P_{(h_1,d)}(z|x) L^i_d(z) \right) \tag{23}$$

$$= \left( \sum_y P_{(h_2,u)}(y|x) L^i_u(y) \right) \left( \sum_y P_{(h_2,v)}(y|x) L^i_v(y) \right) \prod_{d \in D_{uv}} \left( \sum_z P_{(h_1,d)}(z|x) L^i_d(z) \right) \tag{24}$$

$$\text{(because } P_{(h_2,u)} = P_{(h_1,u)}, \text{ and } P_{(h_2,v)} = P_{(h_1,v)}) \tag{25}$$

$$= L^i_{h_2}(x) \prod_{d \in D_{uv}} \left( \sum_z P_{(h_1,d)}(z|x) L^i_d(z) \right) \tag{26}$$

$$= \left( \sum_y P_{(h_2,h_1)}(y|x) L^i_{h_2}(y) \right) \prod_{d \in D_{uv}} \left( \sum_z P_{(h_1,d)}(z|x) L^i_d(z) \right) \tag{27}$$

$$\text{(because } P_{(h_2,h_1)} \text{ is the identity matrix)} \tag{28}$$

It follows that the log-likelihood score remains unchanged subsequent to each transformation operation. The algorithm terminates only if none of the cases apply, which will happen only if $T_\rho$ is a leaf-labeled phylogenetic tree. □

#### 2 Characteristics of simulated data

Characteristics of simulated sequences are presented in Table 1.

#### 3 Results on simulated data

Recall values for subtree size threshold of 10, 20, 30 and 40 are shown in Table 2.

#### 4 Empirical data

Characteristics of empirical data such as length of trimmed alignment, length of complete alignment and results of $\chi^2$ test measuring significant deviation in base composition from average base composition in shown in Table 4.

#### 5 Model selection

The current section describes the model selection framework UNRESTselector that is used to root unrooted trees inferred using MST-backbone(SEM-GM) using CT-HMM that are parameterized in terms of UNREST rate matrices. UNRESTselector iterates over each edge of an unrooted tree $T = (V, E)$ and performs the following tasks. Let the undirected edge $\{u, v\}$ be the edge under consideration. A rooted tree $T_\rho = (V_\rho, E_\rho)$ is constructed by setting $V_\rho$ to $V \cup \{\rho\}$, $E_\rho$ to $E \backslash \{u, v\} \cup \{\{\rho, u\}, \{\rho, v\}\}$, and directing all edges in $E_\rho$ away from $\rho$. A maximum a posteriori (MAP) estimate of ancestral sequence $X^{\text{MAP}}(h)$ is inferred for each hidden vertex $h$ in $V_\rho$ by fitting a GM model using EM. The edge length $t_e$ of each edge $e = (u, v)$ in $E_\rho$ is defined as the Hamming distance between sequences $X^{\text{MAP}}(u)$, and $X^{\text{MAP}}(v)$. The change in base frequency $\Delta_f(u, v)$ for each edge $(u, v)$ in $E_\rho$ is computed as follows: $\Delta_f(u, v) = \sum_{x \in \{A,C,G,T\}} |f_u(x) - f_v(x)|$ where $f_u(x)$ is the fraction of characters in $X^{\text{MAP}}(u)$ that are $x$. Given a base frequency change threshold $\epsilon_f$, a CT-HMM is defined on the basis of rate categories that are assigned to each vertex in $V_\rho$ as follows. The rate category $\rho_{\text{cat}}$ of the root is set to zero. Vertices are visited by performing a preorder tree traversal. Each non-root vertex $c$ that is visited is assigned the rate category of its parent $p$ if $\Delta_f(p, c)$ is not larger than $\epsilon_f$, otherwise $c_{\text{cat}}$ is set to $p_{\text{cat}} + 1$. A distinct rate matrix $Q^i$ is defined for each rate category $i$. The rate category $Q_{(p,c)}$ for each edge $(p, c)$ in $E_\rho$ is defined as $Q^{c_{\text{cat}}}$. The root probability distribution $\pi_\rho$ is defined as the stationary distribution of the rate matrix $Q^{\rho_{\text{cat}}}$. Parameter estimation is performed by optimizing edge lengths, and rate matrices, iteratively until the log-likelihood score converges, using a convergence threshold of 0.01 log-likelihood units. Edge lengths are optimized using Newton-Raphson. The transition matrix $P_e$ for each $e$ in $E_\rho$ is computed as the matrix exponential $P_e = e^{Q_e t_e}$. It is necessary to constrain the elements of $Q_e$ because it is possible to scale $Q_e$ and $t_e$ such that the product $Q_e t_e$ remains unchanged. The rate matrix $Q^i$ for each rate category $i$ is optimized using a simplex method called Nelder-Mead (Nelder and Mead, 1965) subject to the restriction that a non-diagonal element of $Q^i$ was constrained to be one. We restricted element $Q^i(4, 3)$ to be 1. Subsequent to parameter optimization, each rate matrix is normalized by scaling the elements of $Q^i$ such that $\Sigma_x Q^i(x, x)\pi^i(x)$ equals -1, where $\pi^i$ is the stationary distribution of a homogeneous CT-HMM that is parameterized in terms of $Q^i$(Steel, 2016). The root probability $\pi_\rho$ was computed as the stationary distribution corresponding to $Q_\rho$. Threshold $\epsilon_f$ is initially set to the largest observed change in base composition. For each subsequent iteration, $\epsilon_f$ is set to the largest observed change in base composition that is smaller than the value of $\epsilon_f$ for the previous iteration. Model selection is terminated if BIC increases between successive iterations.

The location of the root is determined by selecting the combination of rooted tree and CT-HMM that minimizes BIC.

BIC for UNRESTselector is computed as

$$\text{BIC} = -\log \text{likelihood} + \log(k) \times (11m + |E_\rho|),$$

where $k$ is the number of alignment columns and $m$ is the number of distinct rate matrices.

Models selected by UNRESTselector and IQ-TREE are shown in Table 5.

A comparison of log-likelihood scores and difference in BIC scores obtained using UNRESTselector and IQ-TREE is shown in Table 6.

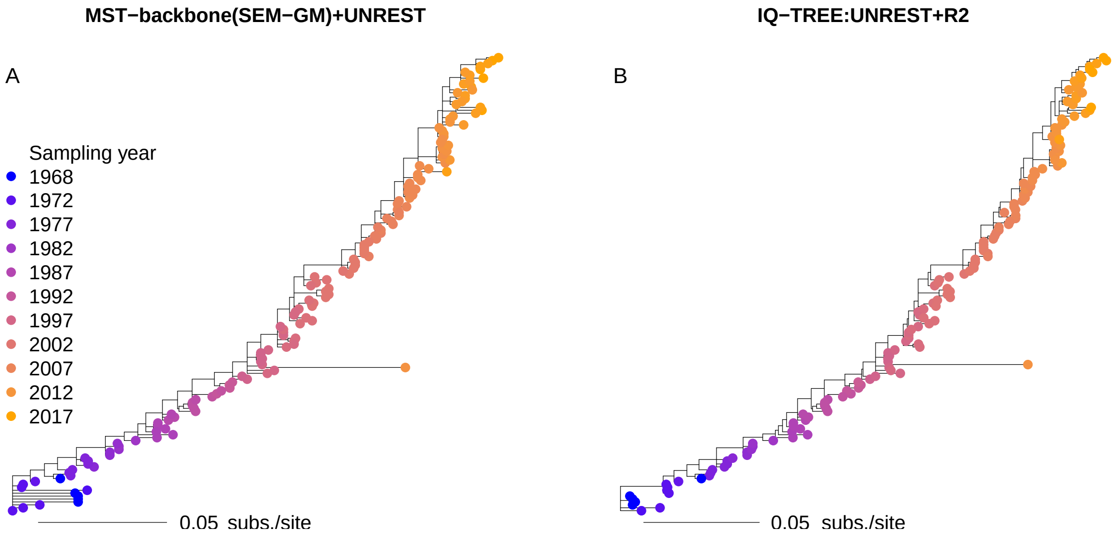


Figure 1: Influenza A H3N2. Bootstrap consensus trees inferred using MST-backbone(SEM-GM) and subsequently rooted under the UNREST model (panel A), and bootstrap consensus trees inferred using IQ-TREE under the UNREST+R2 model. Vertices shown have bootstrap support larger than 70%.

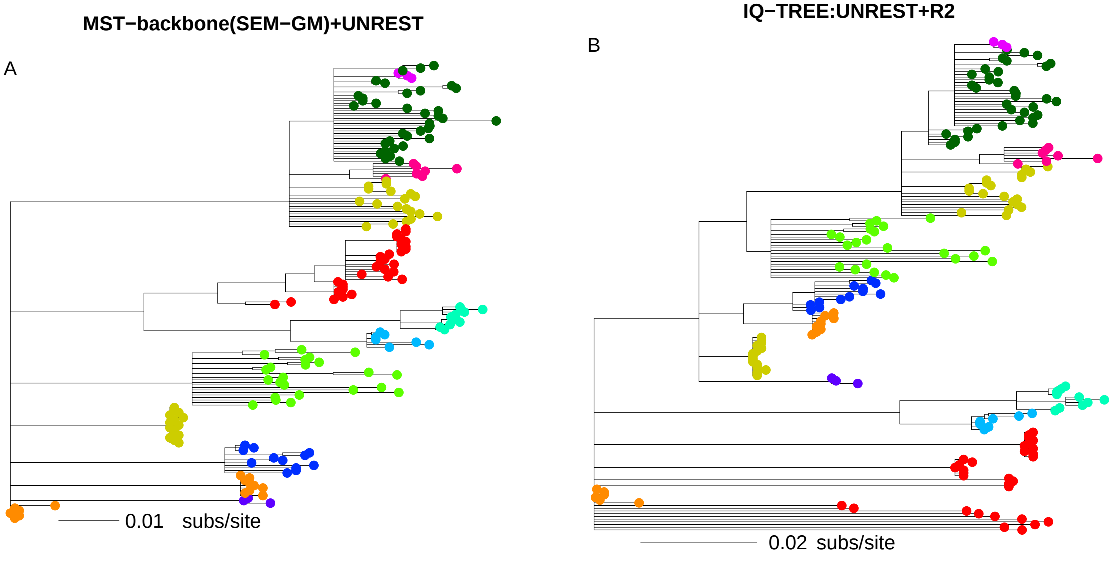


Figure 2: HIV transmission network. Bootstrap consensus trees inferred using MST-backbone(SEM-GM) and subsequently rooted under the UNREST model (panel A), and bootstrap consensus trees inferred using IQ-TREE under the UNREST+R2 model. Vertices shown have bootstrap support larger than 70%

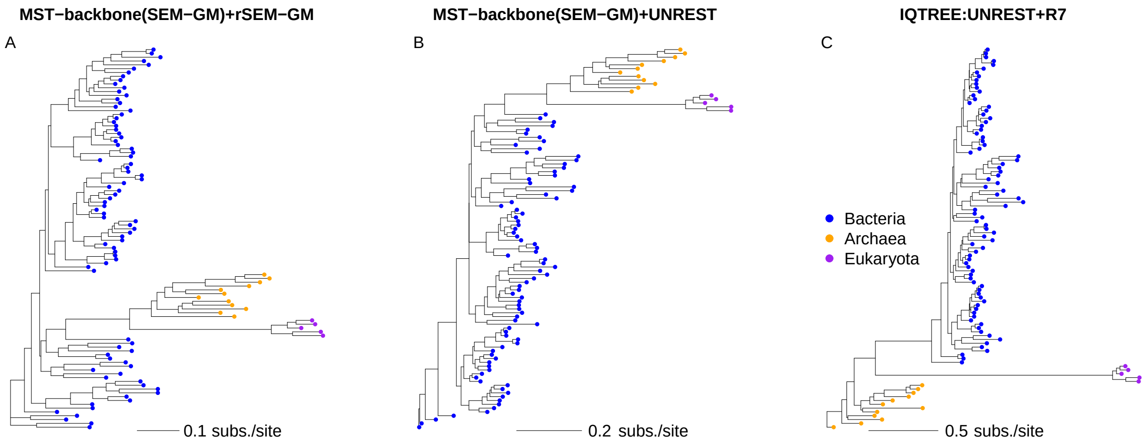


Figure 3: Rooted phylogenetic trees inferred with MST-backbone(SEM-GM)+rSEM-GM (panel A), MST-backbone(SEM-GM)+UNREST (panel B), and IQ-TREE using UNREST+$R_7$(panel C)

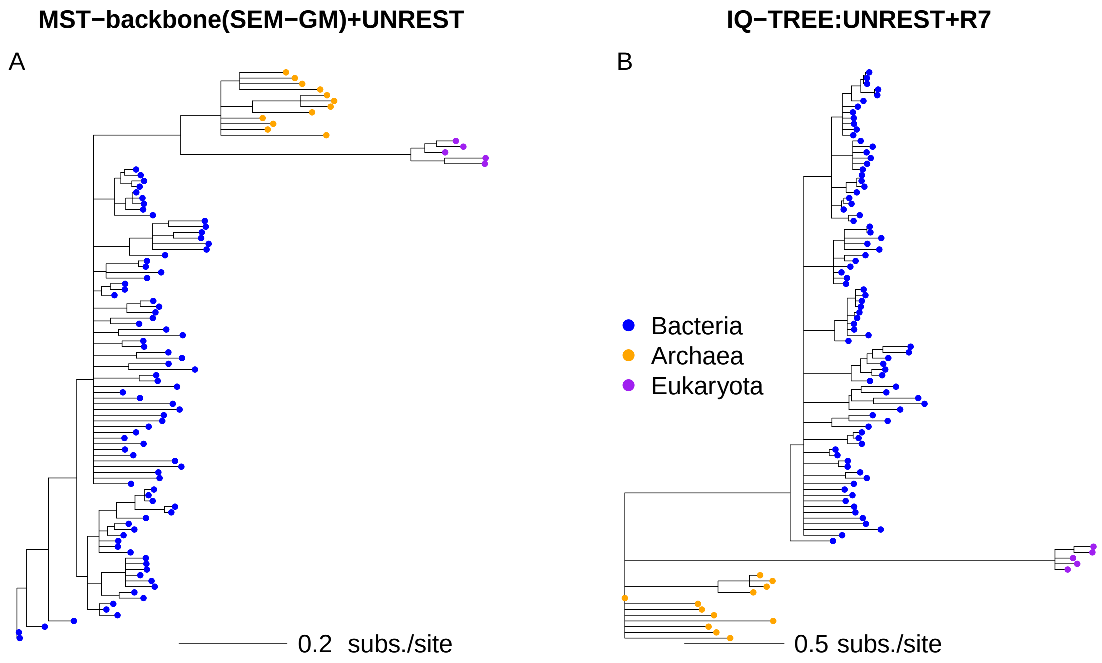


Figure 4: Bootstrap consensus tree for rooted phylogenetic trees inferred with MST-backbone(SEM-GM)+UNREST (panel A), and IQ-TREE using UNREST+$R_7$ (panel B). Vertices shown have bootstrap support larger than 70%

**Algorithm 1:** Transform to bifurcating phylogenetic tree

**Input:** A non-bifurcating tree $T_\rho = (V_{T_\rho}, E_{T_\rho})$, and a GM model $M = (\pi_\rho, \mathbf{P} = \{P_e : e \in E_{T_\rho}\})$

**while** *$T_\rho$ is a non-bifurcating tree* **do**

- **Case** $(i)$**:** $T_\rho$ contains a hidden leaf $h$;
- Let $v$ be the parent of $h$;
- Remove edge $(v, h)$ and matrix $P_{(v,h)}$;
- **Case** $(ii)$**:** $T_\rho$ contain a hidden vertex $h$ with in-degree one and out-degree one;
- Let $u$ and $v$ be the parent and child, respectively, of $h$;
- Remove edges $(u, h)$ and $(h, v)$, and add edge $(u, v)$;
- Remove matrices $P_{(u,h)}$ and $P_{(h,v)}$, and add matrix $P_{(u,v)}$ where $P_{(u,v)} = P_{(u,h)}P_{(h,v)}$;
- **Case** $(iii)$**:** The root is a hidden vertex with out-degree one;
- Let $\pi_\rho^{\text{cur}}$ denote the current root probability distribution;
- Compute new root probability distribution $\pi_\rho^{\text{new}}$ as $\pi_\rho^{\text{new}}(y) = \sum_x \pi_\rho^{\text{cur}}(x)P_{(\rho,v)}(x, y)$, where $v$ is the child of $\rho$;
- Remove edge $(\rho, v)$ and matrix $P_{(\rho,v)}$;
- Set $v$ as the new root of $T_\rho$;
- **Case** $(iv)$**:** $T_\rho$ contains a non-leaf labeled vertex $l$;
- Let $C(l)$ be the children of $l$, and let $h$ be a singleton vertex;
- Add edge $(h, l)$, and add matrix $P_{(h,l)}$, where $P_{(h,l)} = I$ (identity matrix);
- **for** *$v$ in $C(l)$* **do**
  - Remove edge $(l, v)$, and add edge $(h, v)$;
  - Remove matrix and $P_{(l,v)}$, and add matrix $P_{(h,v)}$, where $P_{(h,v)} = P_{(l,v)}$;
- **end**
- **if** *$l$ has a parent $u$* **then**
  - Add edge $(u, h)$ and matrix $P_{(u,h)}$, where $P_{(u,h)} = P_{(u,l)}$;
  - Remove edge $(u, l)$ and matrix $P_{(u,l)}$;
- **else**
  - Set $h$ as the root;
- **end**
- **Case** $(v)$**:** $T_\rho$ contains a hidden vertex $h_1$ with out-degree greater than two;
- Let $u$ and $v$ be a two children of $h_1$ selected at random, and let $h_2$ be a singleton vertex;
- Remove edges $(h_1, u)$ and $(h_1, v)$, and add edges $(h_2, u)$, and $(h_2, v)$, and $(h_2, h_1)$;
- Remove matrices $P_{(h_1,u)}$ and $P_{(h_1,v)}$, and add matrices $P_{(h_2,u)}, P_{(h_2,v)}$, and $P_{(h_2,h_1)}$ such that $P_{(h_2,u)} = P_{(h_1,u)}, P_{(h_2,v)} = P_{(h_1,v)}$, and $P_{(h_2,h_1)} = I$ (identity matrix);

**end**

**Output:** A bifurcating phylogenetic tree $T_\rho$ and a GM model M on $T_\rho$

Table 1: Average edge length and percentage of simulated sequences that reject the null hypothesis of homogeneity in base composition are shown for each setting of $p_{\text{min}}$ that was considered in the main paper. Median and inter-quartile range (shown in parentheses) are listed below.

| $p_{\text{min}}$ | average edge length | $p < 0.05(\%)$ |
|---|---|---|
| 0.7 | 0.15 (0.064) | 89.8 (0.8) |
| 0.8 | 0.1 (0.043) | 83.2 (2.5) |
| 0.9 | 0.05 (0.022) | 68.0 (8.3) |
| 0.95 | 0.025 (0.012) | 53.5 (19.9) |
| 0.99 | 0.005 (0.004) | 19.9 (34.8) |
| 0.995 | 0.0025 (0.0014) | 2.5 (19.1) |

Table 2: Recall values for distinct subtree size threshold

| subtree size | $\text{Re}_S^{\text{nontriv}}$ | $\text{Re}_C^{\text{nontriv}}$ |
|---|---|---|
| 10 | 0.98 (0.0045) | 0.93 (0.035) |
| 20 | 0.98 (0.0055) | 0.93 (0.038) |
| 30 | 0.98 (0.005) | 0.93 (0.038) |
| 40 | 0.98 (0.006) | 0.93 (0.041) |

Table 3: Average branch length for phylogenetic trees inferred using MST-backbone(SEM-GM)

| Gene | average branch length (subs/site) |
|---|---|
| Sanson-all | 0.001 |
| Randall-all | 0.002 |
| Sanson-leaf | 0.002 |
| H3N2 | 0.002 |
| HIV | 0.003 |
| Randall-leaf | 0.03 |
| 16S rRNA | 0.045 |
| COX1 | 0.073 |
| COX3 | 0.084 |
| COX2 | 0.084 |
| ND1 | 0.084 |
| CYTB | 0.087 |
| ATP6 | 0.09 |
| ND4L | 0.095 |
| ND5 | 0.095 |
| ND4 | 0.099 |
| ND3 | 0.106 |
| ATP8 | 0.108 |
| ND6 | 0.116 |
| ND2 | 0.119 |

Table 4: Characteristics of the empirical alignments analyzed in this study

| Gene type | p<0.05 (%) | Original alignment (bp) | Trimmed alignment (bp) |
|---|---|---|---|
| 16S rRNA | 19.3 | 1947 | 971 |
| mt ATP6 | 33.33 | 750 | 652 |
| mt ATP8 | 83.33 | 218 | 128 |
| mt COX1 | 11.11 | 1549 | 1530 |
| mt COX2 | 44.44 | 689 | 659 |
| mt COX3 | 44.44 | 792 | 712 |
| mt CYTB | 11.11 | 1156 | 1085 |
| mt ND1 | 50.0 | 988 | 902 |
| mt ND2 | 5.56 | 1099 | 890 |
| mt ND3 | 50.0 | 361 | 337 |
| mt ND4 | 16.67 | 1391 | 1249 |
| mt ND4L | 77.78 | 306 | 234 |
| mt ND5 | 16.67 | 1773 | 1649 |
| mt ND6 | 38.89 | 561 | 308 |
| Randall-all | 0 | 678 | 678 |
| Sanson-all | 0 | 2236 | 2214 |
| H3N2 | 0 | 1701 | 1701 |
| HIV | 0 | 2873 | 1357 |

Table 5: Results of model selection

| Data | Number of UNREST matrices selected by UNRESTselector | Model selected using IQ-TREE | | |
|---|---|---|---|---|
| | | AIC | AICc | BIC |
| 16S rRNA | 1 | 12.12+R6 | 12.12+R6 | RY8.16+R6 |
| ATP6 | 1 | RY10.12+I+G4 | RY10.12+I+G4 | RY8.18+I+G4 |
| ATP8 | 1 | 12.12+I+G4 | **HKY+F+G4** | **HKY+F+G4** |
| COX1 | 2 | 12.12+I+G4 | 12.12+I+G4 | MK10.34+I+G4 |
| COX2 | 1 | WS10.34+I+G4 | RY8.18+I+G4 | RY8.18+I+G4 |
| COX3 | 1 | WS10.34+I+G4 | WS10.34+I+G4 | WS10.34+I+G4 |
| CYTB | 2 | RY10.12+I+G4 | RY10.12+I+G4 | RY9.20a+I+G4 |
| ND1 | 2 | RY8.10a+I+G4 | RY8.10a+I+G4 | RY8.10a+I+G4 |
| ND2 | 1 | 12.12+I+G4 | 12.12+I+G4 | WS10.34+I+G4 |
| ND3 | 1 | RY9.20a+G4 | RY9.20a+G4 | RY8.18+G4 |
| ND4 | 1 | 12.12+I+G4 | 12.12+I+G4 | WS10.34+I+G4 |
| ND4L | 1 | RY8.18+I+G4 | **TIM+F+I+G4** | **K3Pu+F+G4** |
| ND5 | 1 | WS10.34+I+G4 | WS10.34+I+G4 | RY8.18+I+G4 |
| ND6 | 1 | MK10.34+I+G4 | RY8.18+I+G4 | RY8.18+G4 |
| Sanson-all | 1 | WS5.6a | WS5.6a | **TVMe** |
| Sanson-leaf | 1 | **TVM+F** | **TVM+F** | **TVMe** |
| Randall-all | 1 | WS10.12+R2 | **JC+G4** | **TVMe+G4** |
| Randall-leaf | 1 | WS10.12+I+G4 | WS10.12+I+G4 | WS5.6a+G4 |
| H3N2 | 1 | WS10.12+G4 | WS10.12+G4 | **TVM+F+G4** |
| HIV | 1 | 12.12+R3 | 12.12+R3 | **TVM+F+R3** |

Table 6: Log-likelihood scores and Δ BIC scores of selected models

| Data | log-likelihood | | | Δ BIC | | |
|---|---|---|---|---|---|---|
| | rSEM-GM | UNRESTselector | IQ-TREE | rSEM-GM | UNRESTselector | IQ-TREE |
| Randall-all | -5794.84082 | -4691.58 | -6830.415 | 52064.96 | 0.0 | 4232.03 |
| Randall-leaf | -4367.149902 | -4440.18 | -4446.663 | 2409.52 | 26.14 | 0.0 |
| Sanson-all | -4105.792969 | -3998.85 | -4239.154 | 5235.95 | 0.0 | 418.99 |
| Sanson-leaf | -4094.473389 | -4097.76 | -4215.776 | 2473.64 | 0.0 | 174.41 |
| H3N2 | -10410.4 | -10120.2 | -10992.986 | 25887.79 | 0.0 | 1715.83 |
| HIV | -9908.88 | -9569.18 | -10234.369 | 29185.3 | 0.0 | 1294.31 |
| 16S rRNA | -38999.1 | -40007.96 | -35578.157 | 21726.59 | 8818.35 | 0.0 |
| ND5 | -20720.1 | -21577.82 | -20111.583 | 3943.15 | 2947.29 | 0.0 |
| ND1 | -10352.3 | -10763.84 | -10058.269 | 3092.16 | 1499.6 | 0.0 |
| COX3 | -8040.19 | -8481.22 | -7797.679 | 2888.94 | 1367.09 | 0.0 |
| ATP8 | -1658.93 | -1737.35 | -1664.653 | 1798.36 | 179.36 | 0.0 |
| ND4 | -15833.6 | -16515.7 | -15411.188 | 3454.44 | 2209.03 | 0.0 |
| ND6 | -4202.27 | -4432.22 | -4259.916 | 1999.11 | 361.79 | 0.0 |
| ND4L | -2849.26 | -2987.09 | -2942.106 | 1843.69 | 122.7 | 0.0 |
| COX2 | -7437.28 | -7813.65 | -7246.486 | 2770.18 | 1147.31 | 0.0 |
| ND3 | -4334.61 | -4623.35 | -4322.802 | 2171.22 | 618.55 | 0.0 |
| ND2 | -12739.7 | -13391.26 | -12617.635 | 2729.72 | 1547.26 | 0.0 |
| CYTB | -12562.3 | -12983.66 | -12072.176 | 3545.34 | 1906.84 | 0.0 |
| ATP6 | -7798.17 | -8172.22 | -7582.278 | 2816.44 | 1192.85 | 0.0 |
| COX1 | -15696.6 | -16526.96 | -14925.908 | 4225.27 | 3282.76 | 0.0 |
